# lncRNA ADEPTR loss-of-function elicits sex-specific behavioral and spine deficits

**DOI:** 10.1101/2024.05.25.595903

**Authors:** Kaushik Chanda, Jackson P. Carter, Hirofumi Nishizono, Bindu L Raveendra, Alicia Brantley, Eddie Grinman, Isabel Espadas, Sebastien Lozano-Villada, Jenna Wingfield, Grace Wagner, Amy Peterson, Ryohei Yasuda, Sathyanarayanan V Puthanveettil

## Abstract

Activity-dependent changes in neuronal connections are fundamental to learning and long-term memory storage. However, the precise contribution of long noncoding RNAs (lncRNAs) to these modifications remains unclear. In this study, we assessed the role of the lncRNA ADEPTR, a cAMP-modulated lncRNA localized in dendrites, which is crucial for synapse morphology. By generating two different mouse models—one with a deletion of ADEPTR (L-ADEPTR) and one with a deletion of its protein interaction region (S-ADEPTR)—we investigated the sex-specific impacts of ADEPTR loss of function on learning, memory, dendritic arborization, and synapse morphology. Our behavioral analyses revealed a reduction in anxiety in adult male mice, while learning and memory remained unaffected in both models. Systematic evaluations of neuronal morphology across various developmental stages (∼3-day-old postnatal neuronal cultures and postnatal 14- and 42-day-old male and female mice) uncovered substantial deficits in neuronal architecture in both S- and L-ADEPTR male and female neuronal cultures. At postnatal day 42, in contrast to their male counterparts, L-ADEPTR female mice exhibited a significant deficiency in thin spines. Additionally, we found that the expression of plasticity-related gene BDNF, and immediate early gene cFOS were enhanced in both the cortex and hippocampus of adult male and female S- and L-ADEPTR mice, suggesting the activation of a compensatory mechanism protecting against learning and memory deficits. Collectively, these observations underscore the sex-specific role of lncRNA ADEPTR in shaping neuronal morphology and anxiety behavior.

## Introduction

Dynamic changes in transcriptional networks play a critical role in shaping neuronal connections and their responsiveness to external stimuli, thereby forming the molecular basis of Hebbian plasticity (Ballas et al., 2005; Zheng et al., 2022; Su et al., 2015; Spiegel et al., 2014; Lu et al., 2021). These transcriptional changes are intricately linked to local protein translation (Martin & Ephrussi, 2009; Dahm & Kiebler, 2005; Fernandez-Moya et al., 2014), jointly orchestrating the remodeling of existing synaptic connections and the formation of new ones (Hu et al., 2011; Bailey & Kandel, 2008). Advances in genome and transcriptome sequencing, and sophisticated analyses, have provided a detailed molecular portrait of these transcriptional alterations, unveiling novel classes of noncoding RNAs (ncRNAs) potentially involved in plasticity and memory. Among these, long noncoding RNAs (lncRNAs) have emerged as key players in modulating neuronal maturation, function, memory, and brain pathologies (Andersen et al., 2019; Grinman et al., 2019a; Li et al., 2017; Kim et al., 2023; Chanda et al., 2022; Wei et al., 2022; Ben-Tov Perry et al., 2022; Madugalle et al., 2023; Zou et al., 2023; Chanda et al., 2021; Zhang et al., 2012). Of particular significance are cytoplasmically enriched lncRNAs, which regulate RNA turnover and protein synthesis either directly or by modulating miRNA activity targeting mRNAs (Douka et al., 2021; Kleaveland et al., 2018, Grinman et al., 2019b; Noh et al., 2018; Grinman et al., 2021; Liau et al., 2023, Soutschek and Schratt, 2023; Espadas et al., 2024).

Despite significant advancements, genetic evidence validating the in vivo functions of many long non-coding RNAs (lncRNAs) remains limited. Recent studies have begun to address this gap by generating genetically modified animals with either loss or gain of function in specific lncRNAs (Mattick, 2013; Sauvageau et al., 2013; Ip et al., 2016; Modarresi et al., 2021). Given the large number of expressed lncRNAs, it is urgent to elucidate their functional roles through genetic analyses, especially considering their implications in neurobiological processes such as structural changes, synaptic transmission, transcriptional regulation, and complex behaviors like drug addiction and memory formation. Although genetically modified animals have been successfully developed to study nuclear-enriched lncRNAs and their mechanisms (Zhang et al., 2012; Sauvageau et al., 2013; Ben-Tov Perry et al., 2023), there is a lack of models for investigating cytoplasmically enriched lncRNAs. Recent research highlights the importance of these cytoplasmic lncRNAs in modulating neuronal morphology and memory (Grinman et al., 2021; Cui et al., 2022; Liau et al., 2023; Espadas et al., 2024). Additionally, the potential for sex-specific functions among lncRNAs is an area that remains largely unexplored. A prominent example is XIST (X-inactive specific transcript), a sex-dependent lncRNA predominantly expressed in females, which mediates dosage compensation by silencing the inactive X chromosome (Brown et al., 1991; Lee, 2011).

Sex-specific gene expression patterns have also been documented in various mammalian organs, including the brain (Sarropoulos et al., 2019; Rodríguez-Montes et al., 2023). Recent investigations have delved into the effects of sex-dependent lncRNAs in specific brain regions. For instance, the lncRNA FEDORA, enriched in oligodendrocytes and neurons, was found to be upregulated solely in the prefrontal cortex (PFC) of depressed females (Issler et al., 2022). Overexpression of FEDORA in either neurons or oligodendrocytes exacerbated depression-like behaviors exclusively in female mice, influencing synaptic properties, myelin thickness, and gene networks. Similarly, a study identified female-biased lncRNAs implicated in epigenetic regulation, with the neuronal lncRNA LINC00473 found to modulate synaptic function and gene expression specifically in female mice as a CREB effector (Issler et al., 2020a).

To address these challenges, we carried out genetic analyses of a previously discovered lncRNA, ADEPTR (Activity DEPendently Transported lncRNA), which we identified as a cAMP-regulated gene (Grinman et al., 2021). Remarkably, ADEPTR, residing within the protein-coding gene Arl5b, exhibits upregulation independently of its host gene. Furthermore, ADEPTR localizes to dendrites, facilitated by the molecular motor protein Kif2a. Through our investigation, we pinpointed a 222-nucleotide segment of ADEPTR crucial for its interaction with structural proteins ankyrin (AnkB) and spectrin (Sptn1). Importantly, the loss of ADEPTR function resulted in impaired excitatory synaptic transmission, compromised structural plasticity, and reduced activity-dependent spine enlargement (Grinman et al., 2021). To elucidate its role further, we created two ADEPTR knockouts using CRISPR-Cas9: a long knockout (KO) with the entire ADEPTR gene deleted and a short KO with a specific deletion in ADEPTR’s protein-binding region (from nucleotide position 2415 to 2700).

Using these KO models, we investigated the neurobiological and behavioral consequences of ADEPTR loss of function in both male and female mice. While both male and female KO mice exhibited normal developmental trajectories, male S-ADEPTR mice showed reduced anxiety and L-ADEPTR female mice showed a specific decrease in thin spines. Interestingly, the molecular and neuronal morphology changes observed during early development in both sexes were compensated later in development and into adulthood. These findings underscore the intricate regulation of lncRNAs in modulating neuronal morphology and memory processes.

## Results

### Generation of CRISPR-Cas9 mediated knockout mouse models

To investigate the in vivo functions of ADEPTR, we developed two novel KO mouse models utilizing CRISPR-Cas9 technology. ADEPTR RNA, a single exon transcript, resides within the first intron of its host gene, Arl5b, with specific genomic coordinates outlined in Figure 1A. Using Cas9-mediated editing, we designed two crRNAs targeting either the entire ADEPTR sequence (long deletion, L-ADEPTR) or approximately 200 nucleotides containing protein-binding sites (short deletion, S-ADEPTR), as depicted in Supplementary Figure 1B. The L-ADEPTR KO mouse model was engineered to elucidate the effects of genetic deletion of ADEPTR on developmental processes and brain function, while the S-ADEPTR KO model aimed to delineate the significance of the protein interaction domain in ADEPTR’s in vivo function.

**Figure 1.**
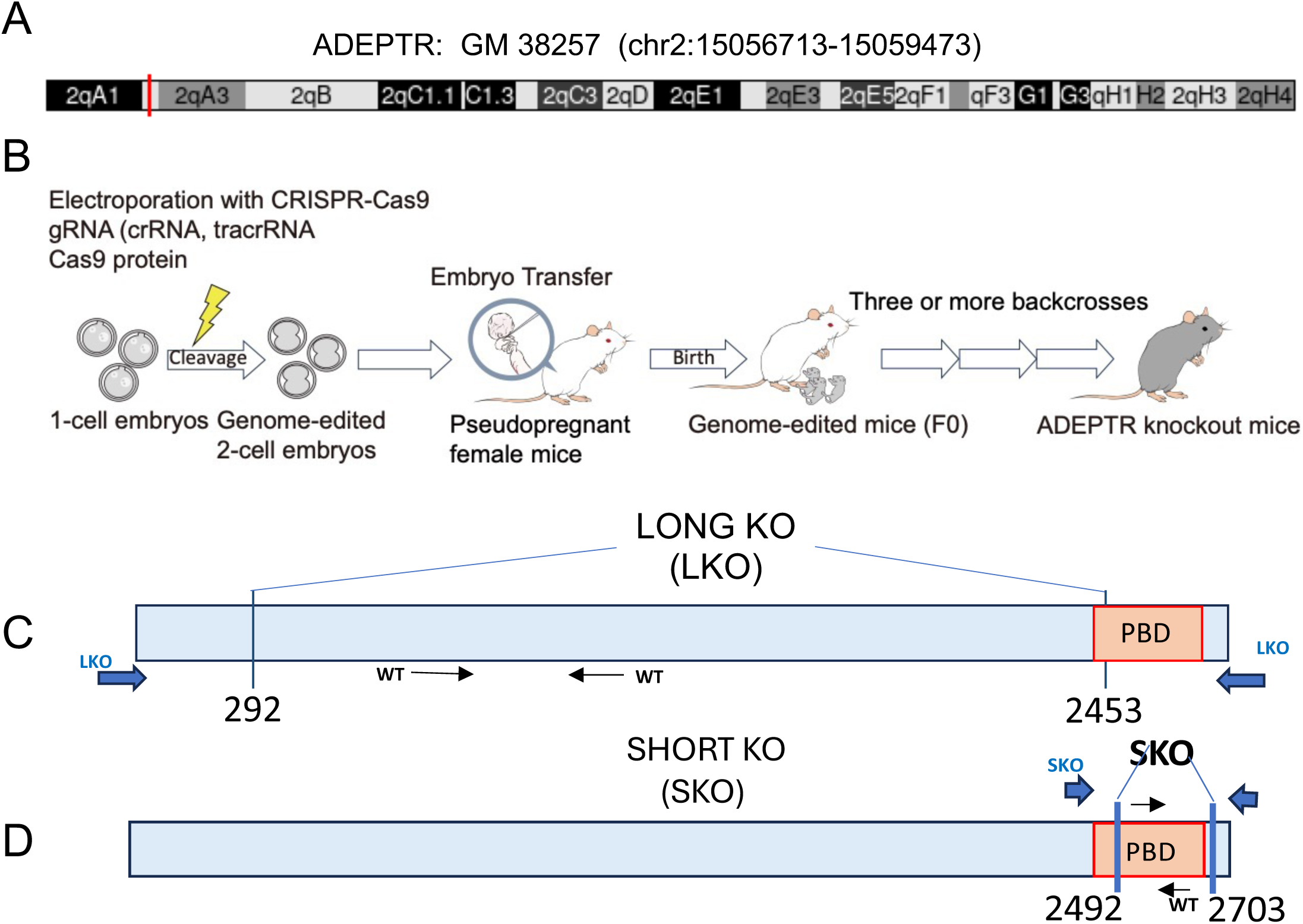
Generation of ADEPTR knockout mice. (A) Chromosomal position of mouse ADEPTR gene. (B) Schematics for the generation of knockout mice by genome editing. (C). Genotyping strategy for Long KO (LKO) ADEPTR mouse (deleted region from 292 bp −2453 bp). The orange region is the Protein binding domain (272bp). Blue thick arrows represent primer binding regions for identifying LKO and black thin arrows indicate primer binding regions for identifying WT. (D) Genotyping strategy of Short Knock out (SKO) ADEPTR (deleted region from 2492 bp −2703 bp). Blue thick arrows represent primer binding regions for identifying SKO and black thin arrows indicate primer binding regions for identifying WT. See Supplementary Figure 1 and Experimental Methods for genotyping details.

Following in vitro annealing of the designed crRNAs with tracrRNA, they were introduced into fertilized eggs of C57BL/6N mice via electroporation alongside Cas9 protein (Figure 1B). Subsequently, the genome-edited fertilized eggs were transferred into pseudo-pregnant mice, and the resultant F0 mice were backcrossed over four generations to establish ADEPTR KO lines.

Two genotyping strategies were employed to identify KO mice. Figure 1C illustrates the primer design schema for validating KOs. To detect wild-type (WT) mice, primer pairs were situated within the Long KO region of ADEPTR. For the Long KO (LKO, position 292bp-2453bp), primer pairs were positioned just outside the two flanking ends of ADEPTR. A representative DNA gel image (Supplementary Figure 1A) demonstrates that this design facilitated robust discrimination between WT and homozygous ADEPTR mice. Specifically, WT mice yielded a band around 500 bp, while the LKO primers generated a band around 650 bp, confirming homozygosity. The absence of bands with the WT primers in homo samples substantiated efficient KO. Additionally, LKO primer pairs failed to amplify in WT mice due to the large size of the amplicon under our PCR conditions. A similar genotyping strategy was employed for SKO (short knockout) ADEPTR mice (deletion of the Protein Binding domain of ADEPTR – 2439-2660bp), as depicted in Fig 1D. The SKO primers were designed to anneal just outside the Protein Binding Domain (PBD) (highlighted in orange), while the WT primers annealed within the PBD (2492-2703bp). SKO mice were genotyped by a commercial source, TransnetyX. Primer sequences for identifying WT, LKO, and SKO are provided in Supplementary Table S1. Remarkably, both L and S ADEPTR KOs did not exhibit embryonic lethality but instead developed normally without any discernible alterations in feeding, growth, or reproduction.

### SKO but not LKO ADEPTR mice display altered anxiety-like behavior

We conducted multiple behavioral analyses on male and female S-ADEPTR mice, alongside wild-type controls housed in their home cages (Fig 2A). To assess anxiety behavior, male and female SKO and LKO mice aged 9-10 weeks underwent an Elevated Plus Maze (EPM) test regimen (Salomons et al., 2012). Our results revealed a notable difference between homo S-ADEPTR females and males in the percentage of time spent in the open arms of the maze (Figure 2B, Supplementary Table S2, female mean - 18.17 sec vs male mean - 31.63, p-value - 0.032, n = 16 per group, one-way ANOVA). Conversely, there was no disparity between WT S-ADEPTR females and males (female mean - 18.67 sec vs male mean - 22.45, n = 16 per group, one-way ANOVA followed by Tukey’s test).

**Figure 2:**
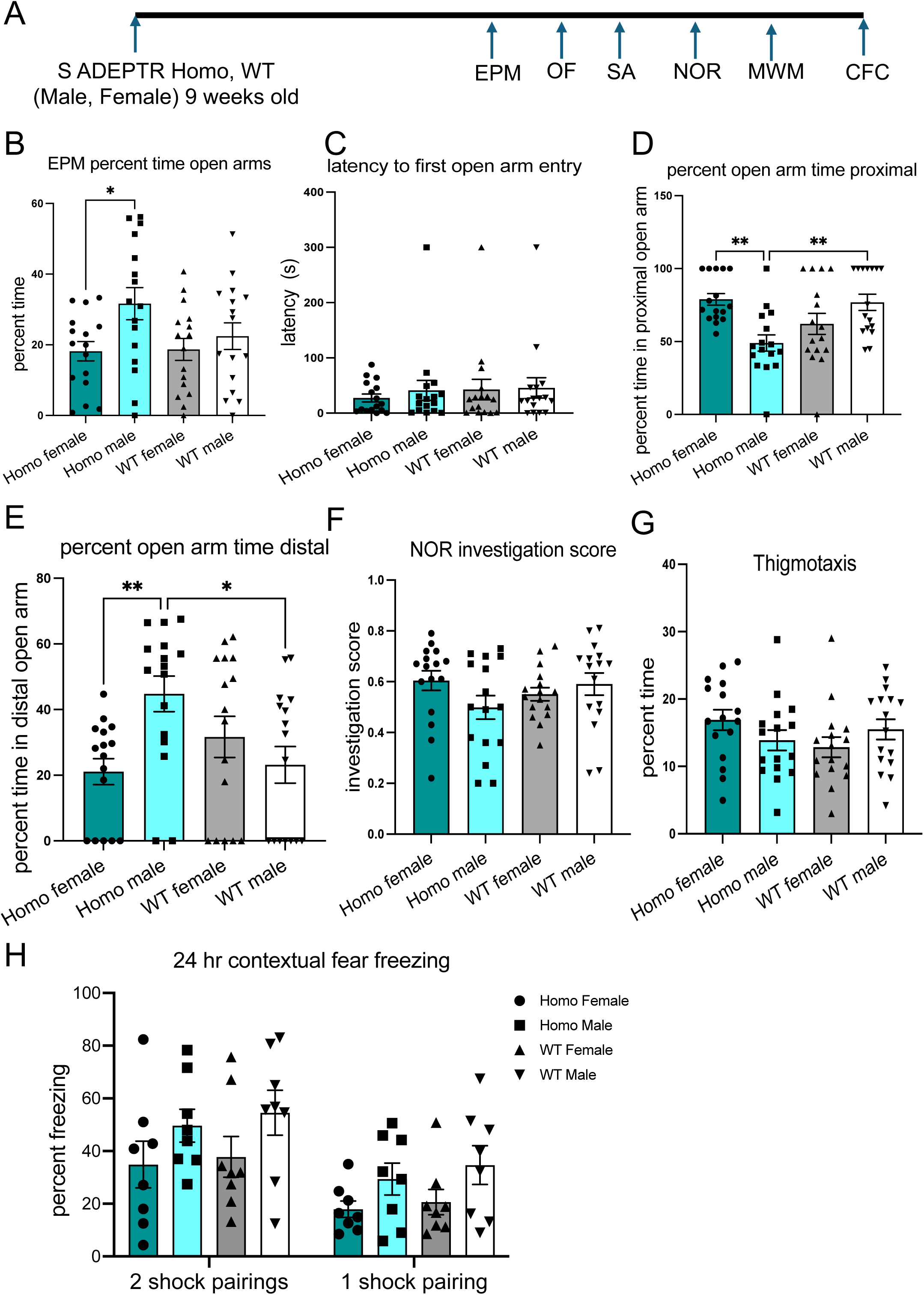
SKO mice display altered anxiety-like behavior but no deficits in memory. (A) Experimental strategy to assess behavior in S-ADEPTR Homozygous (Homo) and Wild-type (WT) controls. (B) Bar Graph depicting percent time spent in open arms in Elevated Plus Maze (EPM) of indicated phenotypes (N=14-15 per group) of S-ADEPTR mice. (C) Latency (s) in Elevated Plus Maze of indicated phenotypes (N=14-15 per group). (D) Percent time spent in proximal open arms in Elevated Plus Maze of indicated phenotypes (N=14-15 per group). (E) Percent time spent in distal open arms in Elevated Plus Maze of indicated phenotypes (N=14-15 per group). (F) Investigation score of Novel object recognition (NOR) in indicated phenotypes (N=15-16 per group). (G) Percent time (s) in Thigmotaxis of indicated phenotypes (N=15-16 per group), one way ANOVA followed by Tukey’s test. (H) Percent freezing (s) in 1 or 2 shock pairings in 24-hour Contextual fear conditioning (CFC) of indicated phenotypes (N=8-9 per group). Error bars indicate ± SEM. Significance level between different experimental pairs is shown (NS, not significant; \**p* < 0.05; \*\**p* < 0.01; \*\*\**p* < 0.001). See Supplementary Figure 1 for Open Field (OF), Spontaneous Alternation (SA), and Morris Water Maze (MWM) assessments and Supplementary Figure 2 for Reversal Learning in MWM.

Furthermore, we found no significant variance in the latency to the first open arm entry between WT and homo S-ADEPTR mice (for both males and females) (Figure 2C, n = 16 per group, one-way ANOVA). However, homo S-ADEPTR male mice exhibited a reduced duration in the proximal open arm compared to homo S-ADEPTR females (Figure 2D, female mean - 78.91 sec vs male mean - 48.98, p-value - 0.0012, one-way ANOVA). Upon closer examination, we identified significant differences between WT S- ADEPTR males and homo S-ADEPTR males (WT male mean - 76.84 sec vs homo male mean - 48.98, p-value - 0.0030, one-way ANOVA), while no such distinction was observed between WT females and homo S-ADEPTR females. Additionally, homo S- ADEPTR males spent more time on the distal arm compared to homo S-ADEPTR females (Figure 2E, homo female mean - 21.08 sec vs homo male mean – 44.76 sec, p-value - 0.0084, n = 16 per group, one-way ANOVA) or WT males (WT male mean - 23.15 sec vs homo male mean – 44.76 sec, p-value - 0.0182).

Collectively, these findings suggest that male homo S-ADEPTR male mice exhibit significantly lower anxiety-like behavior and potentially higher risk-taking compared to female homo S-ADEPTR mice. However, such a trend was not evident in the EPM test conducted on LKO mice with a similar experimental strategy (Figure 3A-E, Supplementary Table S3, n = 13-15 per group, 2-way ANOVA followed by Tukey’s test).

**Figure 3:**
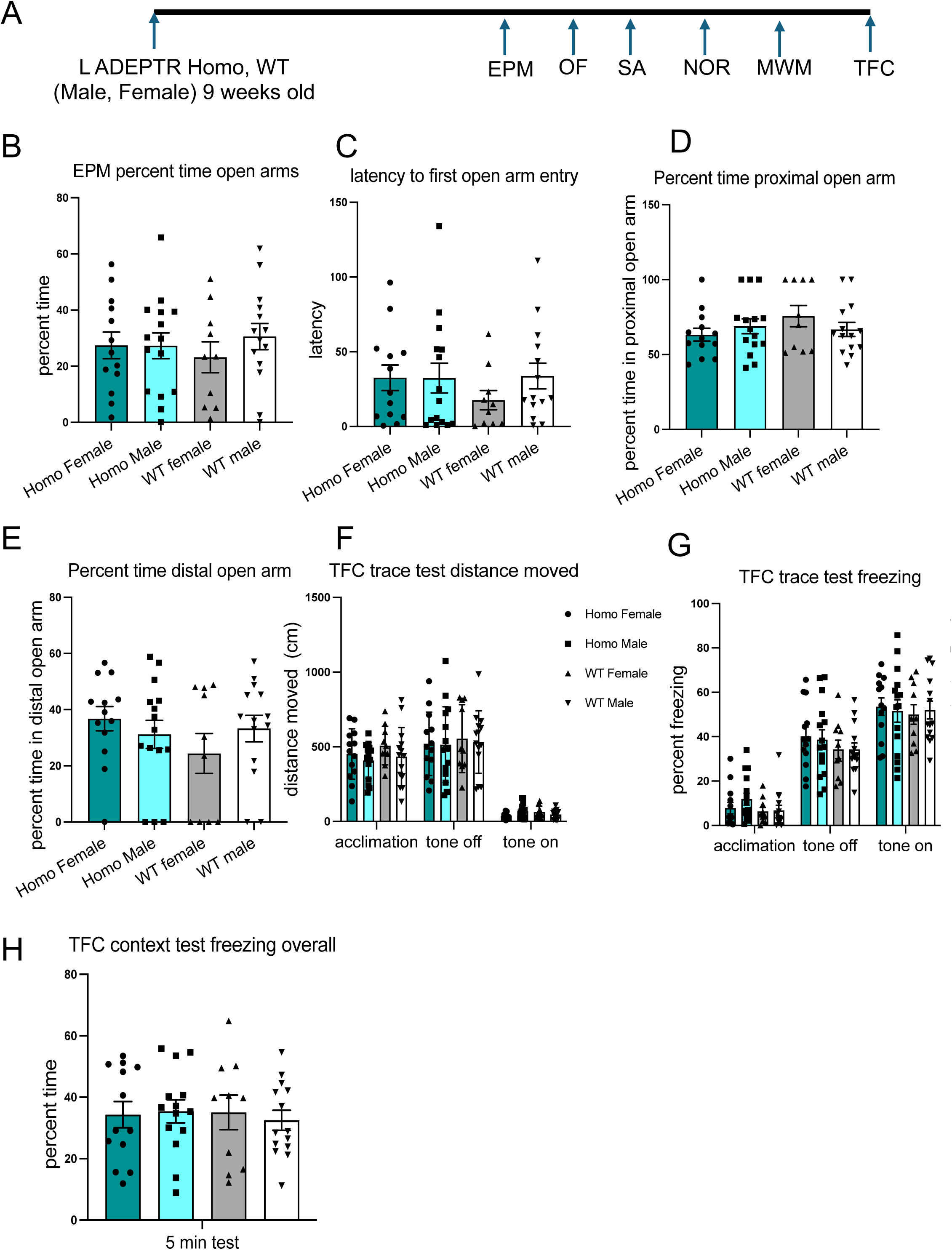
LKO mice display no deficits in anxiety-like behavior or memory. (A) Experimental strategy to access behavior in L-ADEPTR Homozygous (Homo) mice and Wild-Type (WT) controls. (B) Bar Graph depicting percent time spent in open arms in Elevated Plus Maze (EPM) of indicated phenotypes (N=13-15 per group) of L-ADEPTR mice. (C) Latency (s) in Elevated Plus Maze of indicated phenotypes (N=13-15 per group). (D) Percent time spent in proximal open arms in Elevated Plus Maze of indicated phenotypes (N=13-15 per group). (E) Percent time spent in distal open arms in Elevated Plus Maze of indicated phenotypes (N=13-15 per group). (F) Distance moved (cm) under acclimation, with or without tone in Trace Fear conditioning (TFC) of indicated phenotypes (N=13-15 per group). (G) Percent freezing (s) under acclimation, with or without tone in Trace Fear conditioning of indicated phenotypes (N=13-15 per group). (H) Percent time (s) test freezing overall in Trace Fear conditioning of indicated phenotypes (N=13-15 per group), One way ANOVA followed by Tukey’s test. Error bars indicate ± SEM. Significance level between different experimental pairs is shown (NS, not significant; \**p* < 0.05; \*\**p* < 0.01; \*\*\**p* < 0.001). See Supplementary Figure 3 for Novel Object Recognition (NOR), Open Field (OF), Spontaneous Alternation (SA), and Morris Water Maze (MWM) assessments and Supplementary Figure 4 for reversal Learning in MWM.

### SKO and LKO ADEPTR mice show no deficits in memory

Our prior research has demonstrated synaptic targeting of ADEPTR in a cAMP-dependent manner, independent of its protein-coding host gene (Grinman et al., 2021). Additionally, ADEPTR exhibits specific regulation by excitatory signaling, with its cAMP-induced expression relying on PKA activity. Unlike Arl5b, ADEPTR expression is negatively regulated by GABA exposure. Building upon this foundation, we proceeded to evaluate memory formation in SKO and LKO mice using a battery of tests, including Spontaneous Alternation (for working memory), Novel Object Recognition (for non-spatial memory), Morris Water Maze (MWM) (for spatial memory and cognitive flexibility), and Contextual Fear Conditioning (for memory acquisition and consolidation).

In the Novel Object Recognition (NOR) test, no significant difference in investigation score was observed between WT and homozygous (for both male and female S-ADEPTR) mice, with scores above the 50% chance level (Figure 2F, Supplementary Table S2, n = 16 per group, one-way ANOVA). Similarly, in the Open Field (OF) test, there were no significant differences in activity during 10-minute intervals between WT and homozygous mice (for both male and female) (Figure 2G and Supplementary Fig 1C, n = 16 per group, one-way ANOVA). Furthermore, no significant differences were detected in spontaneous alternation (SA) between WT and homozygous mice (for both male and female), with or without delay (Supplementary Figure 1D-E, Supplementary Table S2, n = 16 per group, one-way ANOVA). Similarly, in the Morris water maze (MWM), no significant differences were observed between WT and homozygous mice (for both male and female) across the six parameters tested (Supplementary Figure 1F-K and Supplementary Fig 2 A-F, n = 16 per group, one-way ANOVA). Contextual Fear Conditioning (CFC) revealed no significant difference in freezing percent between WT and homozygous mice (for both male and female) under both 1-shock and 2-shock pairings (Figure 2H, n = 8 per group, one-way ANOVA).

Interestingly, male and female L-ADEPTR mice also exhibited no significant differences in these memory assessments. In the NOR assay, investigation scores did not significantly differ between WT and homozygous mice (for both male and female L-ADEPTR), exceeding the 50% chance level (Supplementary Figure 3A, Supplementary Table S3, n = 10-15 per group, one-way ANOVA). Similarly, there were no significant differences in activity during 10-minute intervals in the OF test or thigmotaxis between WT and homozygous mice (for both male and female L-ADEPTR) (Supplementary Figure 3B, n = 13-15 per group, one-way ANOVA), nor in the SA test (Supplementary Figure 3C-D, n = 13-15 per group, one-way ANOVA), nor in MWM learning (Supplementary Figure 3E-H and Supplementary Fig 4A-D, n = 13-15 per group, one-way ANOVA). In the L-ADEPTR mice, the Trace Fear conditioning (TFC) paradigm revealed no significant differences between WT and homozygous mice (for both male and female L-ADEPTR) during acclimation, and with or without tone, across the three parameters tested – trace test distance moved, trace test freezing, and context test freezing (Figure 3F-H, Supplementary Table S3, n = 13-15 per group, 2-way ANOVA).

### ADEPTR loss-of-function impairs dendritic arbor and spine density in both sexes

The absence of a phenotype in our memory assessment paradigms prompted an investigation into whether S- or L-ADEPTR KOs exhibit deficiencies in spine morphology. Drawing from our previous findings in neuronal cultures, which demonstrated that RNAi-mediated loss of ADEPTR function resulted in reduced spine density and mushroom spines (Grinman et al., 2021), we anticipated similar deficiencies in spine density and morphology in ADEPTR KO mice. Given the established association between spine density and morphology changes and learning in numerous studies, the lack of a memory phenotype suggested potential stability in spine density and/or morphology in ADEPTR KO mice.

To explore these possibilities, we initially examined the neuronal morphology of primary cultured neurons derived from male and female ADEPTR KO mice. Specifically, we analyzed dendritic arborization, spine density, and spine morphology in primary hippocampal neurons from both homozygous female and male mice (P1-P3) compared to WT controls. Our results revealed a significant reduction in distal branches (70-110 μm) in homozygous female S-ADEPTR mice compared to WT littermate controls, along with a mild reduction in distal secondary branches (40-60 μm), while no differences were observed in proximal secondary branches (20-40 μm) (Figure 4A, B, Supplementary Table S4, WT, n=16, Homo Female, n=11 paired t test, *p<0.05, **p<0.005). In Homo S-ADEPTR males, there was significant reduction in branching uniformly in the secondary as well as tertiary branching (Figure 4C, D, WT, n=19, Homo Male, n=9 paired t test, *p<0.05, **p<0.005). Similarly, L-ADEPTR mice showed significant reduction in arborization in the distal secondary and tertiary branching in Homo L-ADEPTR males (Figure 5A, B, Supplementary Table S5, WT, n=10, Homo Male, n=14 paired t test, *p<0.05, **p<0.005), as well as Homo L-ADEPTR females (Figure 5C, D, WT, n=12, Homo Female, n=15 paired t test, *p<0.05, **p<0.005).

**Figure 4:**
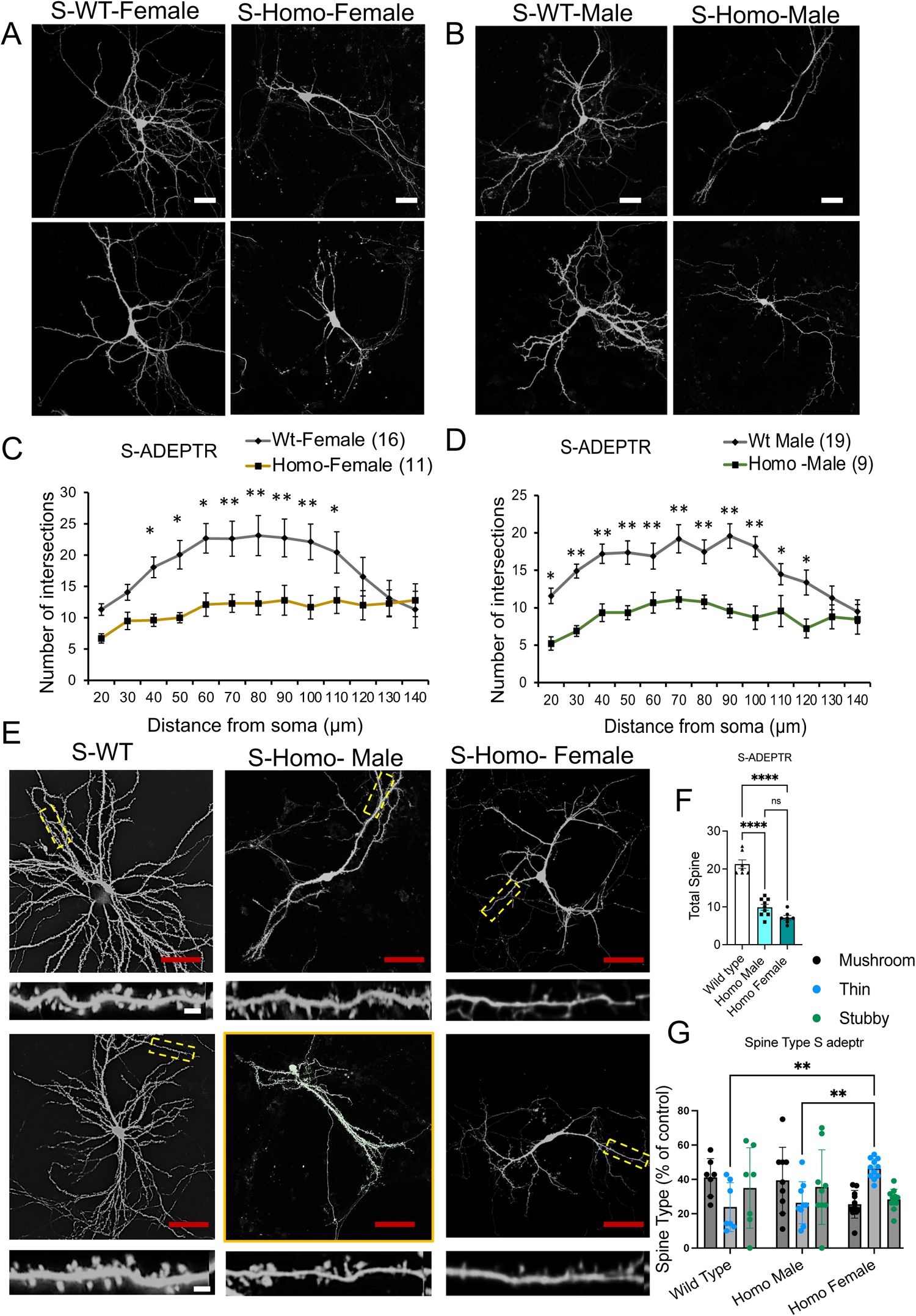
ADEPTR loss-of-function impairs dendritic arbor and spine density in S-ADEPTR mice sex specifically. (A) Representative confocal projection images showing the soma and dendritic arbor of Wild-Type (WT) littermate of S-ADEPTR (S-WT-Female) and Homozygous (Homo) female (S-Homo) mice. Scale Bar=40µm. (B) Representative confocal projection images showing the soma and dendritic arbor of Wild-Type (WT) littermate of S-ADEPTR male (S-WT-Male) and Homozygous (Homo) male (S-Homo) mice. Scale Bar=40µm. (C) and (D) Quantification of dendritic morphology using Sholl analysis of intersections per 10-µm step size. Changes compared between WT and Homo. Paired t-test. Error bars represent SEM. *p<0.05, **p<0.005, ***p<0.0005. (E) Confocal projection images showing area analyzed for spine morphology and enlarged image in the inset for spine details of indicated phenotypes. Scale Bar=40µm. Dendrite inset Scale bar=2µm. (F) Bar graph depicting total spines in WT, Homo male and Homo Female in S-ADEPTR mice, one way ANOVA followed by Tukey’s test (N=9-11 neurons per group). G) Bar graph showing Mushroom, Thin and Subby spine types in WT, Homo male and Homo Female in S-ADEPTR mice (N=7-9 neurons per group). One way ANOVA followed by Tukey’s test. Error bars represent SEM. *p<0.05, **p<0.005, ***p<0.0005.

**Figure 5:**
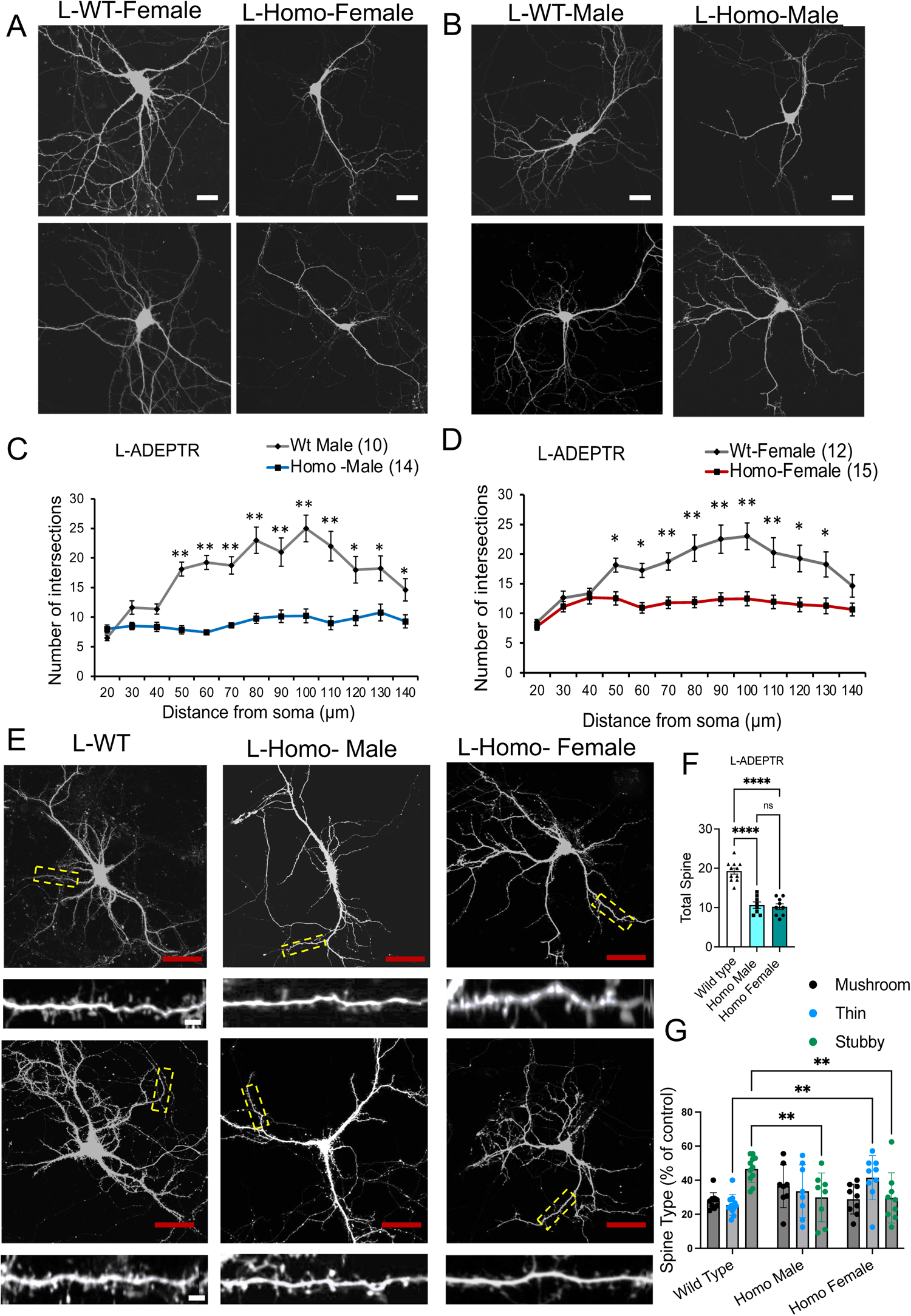
ADEPTR loss-of-function impairs dendritic arbor and spine density in L-ADEPTR mice. (A) Representative confocal projection images showing the soma and dendritic arbor of Wild-Type (WT) littermate of L-ADEPTR Female (L-WT-Female), and Homozygous (L-Homo-female) mice. Scale Bar=40µm. (B) Representative confocal projection images showing the soma and dendritic arbor of Wild-Type (WT) littermate of L-ADEPTR male mice (L-WT-Male), and Homozygous (L-Homo) male mice. Scale Bar=40µm (C) and (D) Quantification of dendritic morphology changes using Sholl analysis of intersections per 10-µm step size. Changes compared between WT and Homo. Paired t-test. Error bars represent SEM. *p<0.05, **p<0.005, ***p<0.0005. (E) Confocal projection images showing area analyzed for spine morphology and enlarged image in the inset for spine details of indicated phenotypes. Scale Bar=40µm. Dendrite inset Scale bar=2µm. (F) Bar graph depicting total spines in WT, Homo male and Homo Female in L-ADEPTR mice, one way ANOVA followed by Tukey’s test (N=9-11 neurons per group). (G) Bar graph showing Mushroom, Thin and Subby spine types in WT, Homo male and Homo Female in L-ADEPTR mice (N=8-10 neurons per group). One way ANOVA followed by Tukey’s test. Error bars represent SEM. *p<0.05, **p<0.005, ***p<0.0005.

A more detailed examination unveiled significant alterations not only in arborization but also in spine density among S-ADEPTR neurons. Specifically, there was a substantial reduction in total spines in homozygous males and S-ADEPTR females compared to their wild-type counterparts (Figure 4E, F, Supplementary Table S4, WT. mean-21.28 ± 1.18 vs Homo male mean - 9.88 ± 1.26 vs Homo Female mean – 7.14 ± 1.22, n=7-9 per group, one-way ANOVA, *p < 0.05, **p < 0.005, ***p < 0.0005, ****p < 0.0005), although no significant differences were observed between homozygous males and females. Analysis of spine morphology distribution indicated no significant differences in mushroom, thin, or stubby spines in homozygous S-ADEPTR males compared to WT controls. However, significant differences were found in the thin spine type between WT and homozygous S-ADEPTR females, as well as between homozygous S-ADEPTR males and females (Figure 4G, n=7-11 per group, one-way ANOVA, Tukey’s test, *p < 0.05, **p < 0.005).

Similarly, homozygous L-ADEPTR males and females exhibited a significant reduction in total spines compared to their wild-type counterparts (Figure 5E, F, Supplementary Table S5, WT. mean-19.27 ± 1.09 vs Homo male mean - 10.62 ± 1.06 vs Homo Female mean – 10.22 ± 1.14, n=8-11 per group, one-way ANOVA, *p < 0.05, **p < 0.005, ***p < 0.0005, ****p < 0.0005). Analysis of spine morphology distribution revealed significant differences in stubby spines in homozygous L-ADEPTR males and females compared to WT controls. Additionally, significant disparities were observed in the thin spine type between WT and homozygous L-ADEPTR females, as well as between homozygous L-ADEPTR males and females (Figure 5G, n=7-11 per group, one-way ANOVA followed by Tukey’s multiple comparisons, *p < 0.05, **p < 0.005).

Taken together, these analyses suggest more severe deficits in neuronal morphology in S-ADEPTR and L-ADEPTR mice compared to RNAi-mediated ADEPTR loss of function in hippocampal neuronal cultures.

### Golgi Staining reveals female specific deficiency in the density of thin spines in hippocampal neurons

Our observation of significant changes in neuronal morphology in both sexes of S-ADEPTR and L-ADEPTR primary neuron cultures, coupled with the absence of significant memory impairments, led us to inquire whether these deficits might be compensated for in adults, resulting in the lack of memory deficits. To investigate this possibility, we examined the brains of male and female mice at two different ages: P14, corresponding to postnatal development, and P42, corresponding to adulthood. Specifically, we conducted Golgi staining of the hippocampus and imaged apical and basal dendrites of male and female S-ADEPTR and L-ADEPTR mice. Quantitative analysis of imaging data revealed that at P14, neither S-ADEPTR (Fig 6 A-J, Supplementary Table S6) nor L-ADEPTR (Fig 7 A-J, Supplementary Table S7) mice exhibited significant changes in dendritic arborization, length, or spine density in both males and females. Moreover, there was no significant reduction in the number of dendritic branch points, as evidenced by sholl analysis.

**Figure 6:**
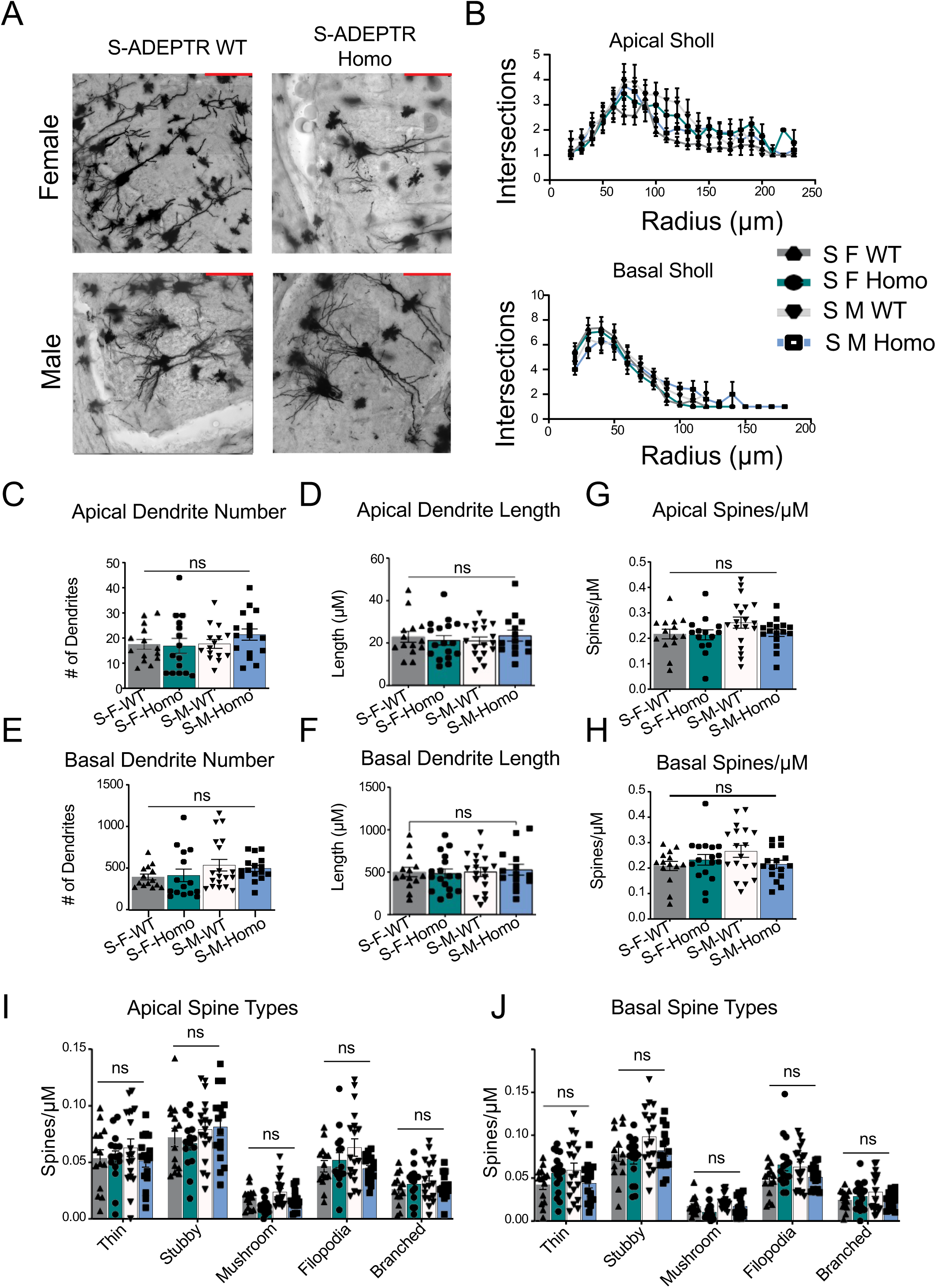
Golgi Staining of P14 S-ADEPTR CA1 pyramidal Hippocampal mouse neurons. (A) Representative images of Golgi stained CA1 pyramidal cells (male (M) and female (F) Wild-type littermate and homozygous (Homo) S-ADEPTR mice. Scale: 100uM. (B) Sholl analysis of P14 S-ADEPTR cells. (C, E) Total number of apical and basal dendrites, respectively. (D, F) Total length of apical and basal dendrites, respectively. (G, H) Spine density of apical and basal dendrites, respectively. (I, J) Spine density of different spine classes on apical and basal dendrites, respectively. (N=14-15 per group). One way ANOVA followed by Tukey’s test. Error bars indicate ± SEM. Significance level between different experimental pairs is shown (NS, not significant; \**p* < 0.05; \*\**p* < 0.01; \*\*\**p* < 0.001).

**Figure 7:**
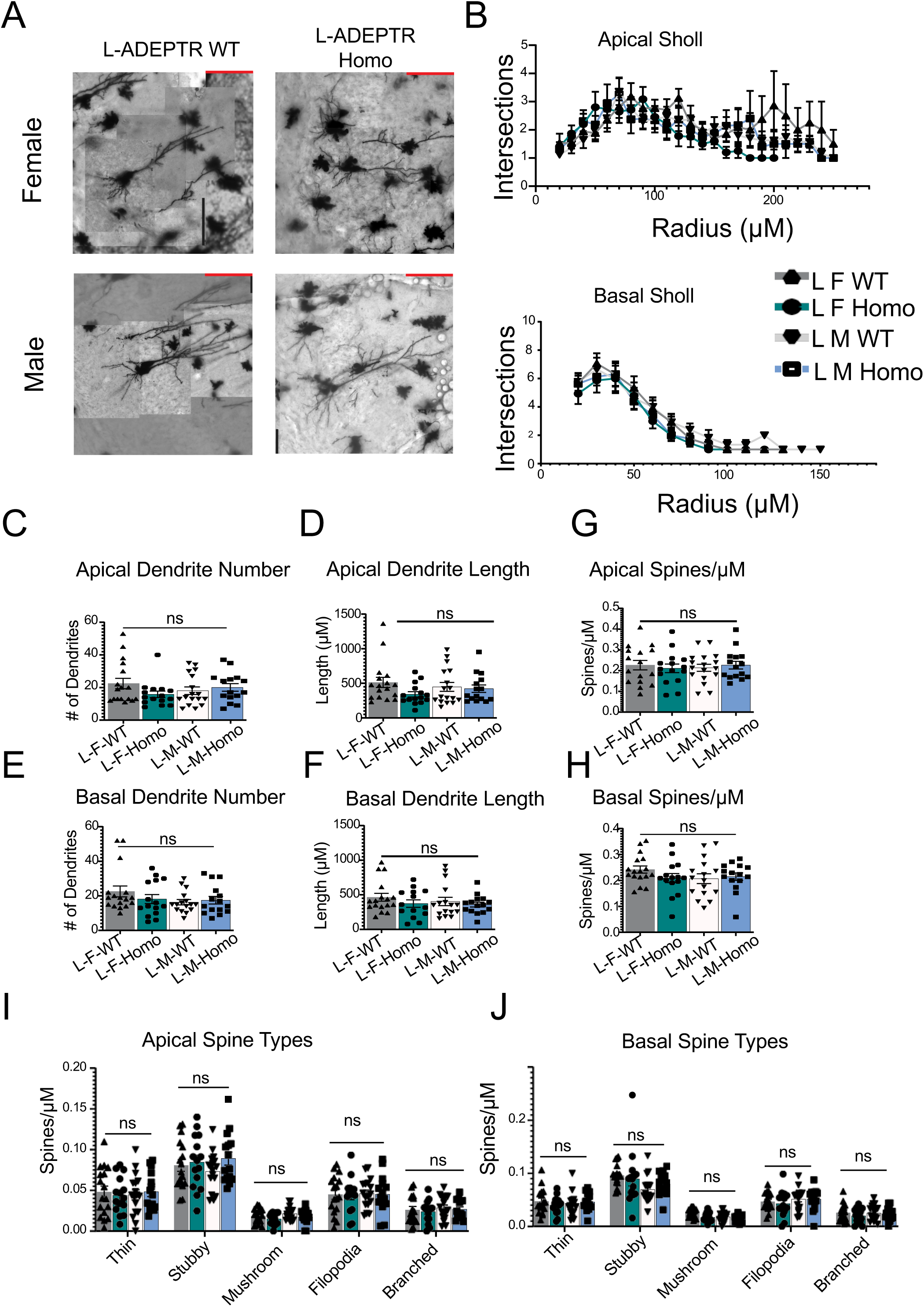
Golgi Staining of P14 L-ADEPTR CA1 pyramidal Hippocampal mouse neurons. (A) Representative images of Golgi stained CA1 pyramidal cells (male (M) and female (F) Wild-type (WT) littermate and homozygous (Homo) L-ADEPTR mice. Scale: 100uM. (B) Sholl analysis of P14 L-ADEPTR cells. (C, E) Total number of apical and basal dendrites, respectively. (D, F) Total length of apical and basal dendrites, respectively. (G, H) Spine density of apical and basal dendrites, respectively. (I, J) Spine density of different spine classes on apical and basal dendrites, respectively. (N=14-15 per group). One way ANOVA followed by Tukey’s test. Error bars indicate ± SEM. Significance level between different experimental pairs is shown (NS, not significant; \**p* < 0.05; \*\**p* < 0.01; \*\*\**p* < 0.001).

However, at P42, a mild morphological phenotype became apparent. Although no changes were observed in S-ADEPTR (Figure 8 A-J, Supplementary Table S8), a modest but significant reduction in thin spines was exclusively observed in L-ADEPTR females (Fig 9J) (p=0.0025, two-way ANOVA), subsequently resulting in a reduction in the total spine density of L-ADEPTR KO females (p=0.0009, two-way ANOVA). No other spine type, nor dendritic length, number, or branching, was affected in S-ADEPTR or L-ADEPTR males or females (Fig 9 A-I, Supplementary Table S9).

**Figure 8:**
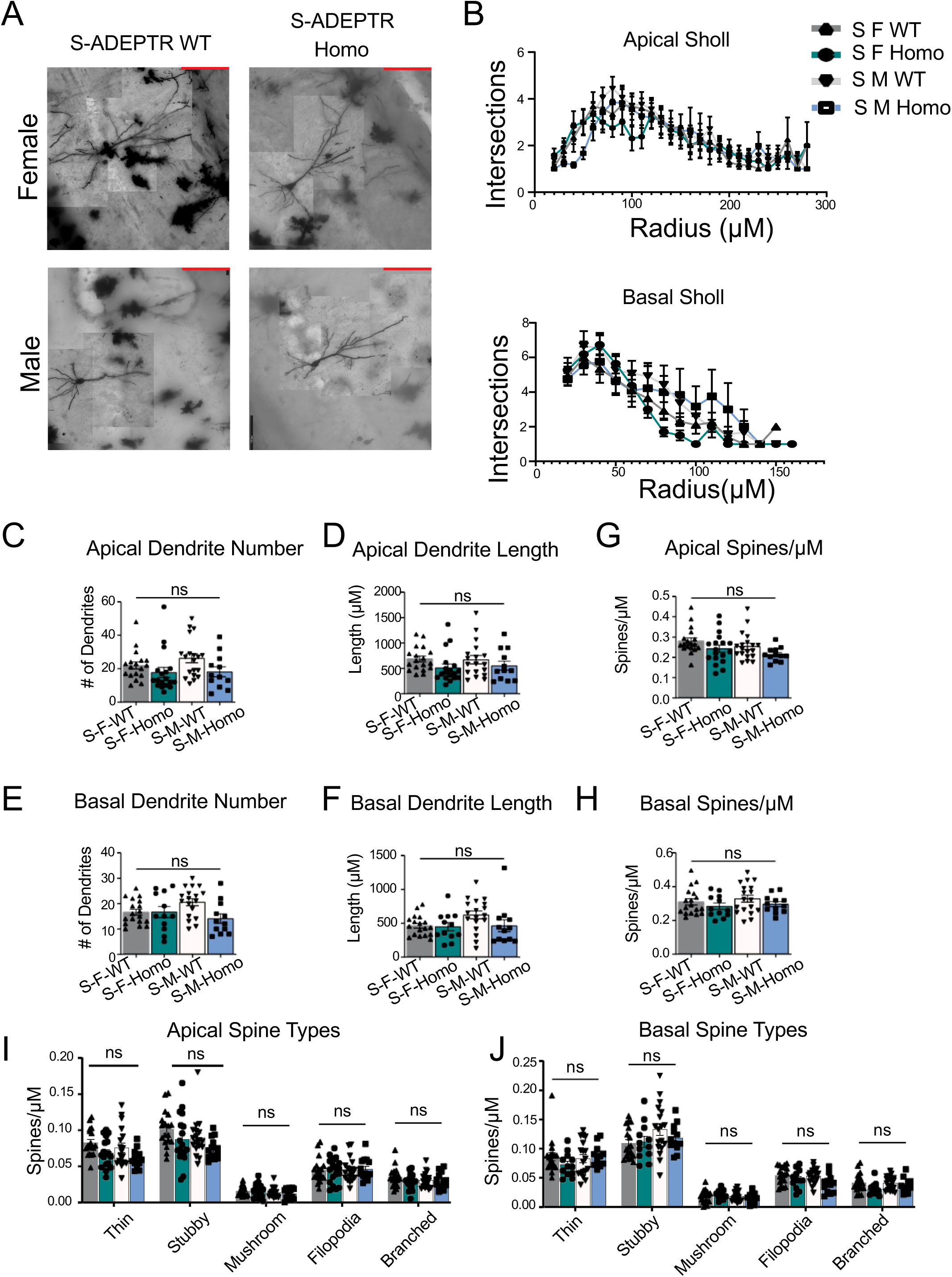
Golgi Staining of P42 S-ADEPTR CA1 pyramidal Hippocampal mouse neurons. (A) Representative images of Golgi stained CA1 pyramidal cells (male (M) and female (F) Wild-type (WT) littermate and homozygous (Homo) S-ADEPTR mice. Scale: 100uM. (B) Sholl analysis of P42 S-ADEPTR cells. (C, E) Total number of apical and basal dendrites, respectively. (D, F) Total length of apical and basal dendrites, respectively. (G, H) Spine density of apical and basal dendrites, respectively. (I, J) Spine density of different spine classes on apical and basal dendrites, respectively. (N=14-15 per group). One way ANOVA followed by Tukey’s test. Error bars indicate ± SEM. Significance level between different experimental pairs is shown (NS, not significant; \**p* < 0.05; \*\**p* < 0.01; \*\*\**p* < 0.001).

**Figure 9:**
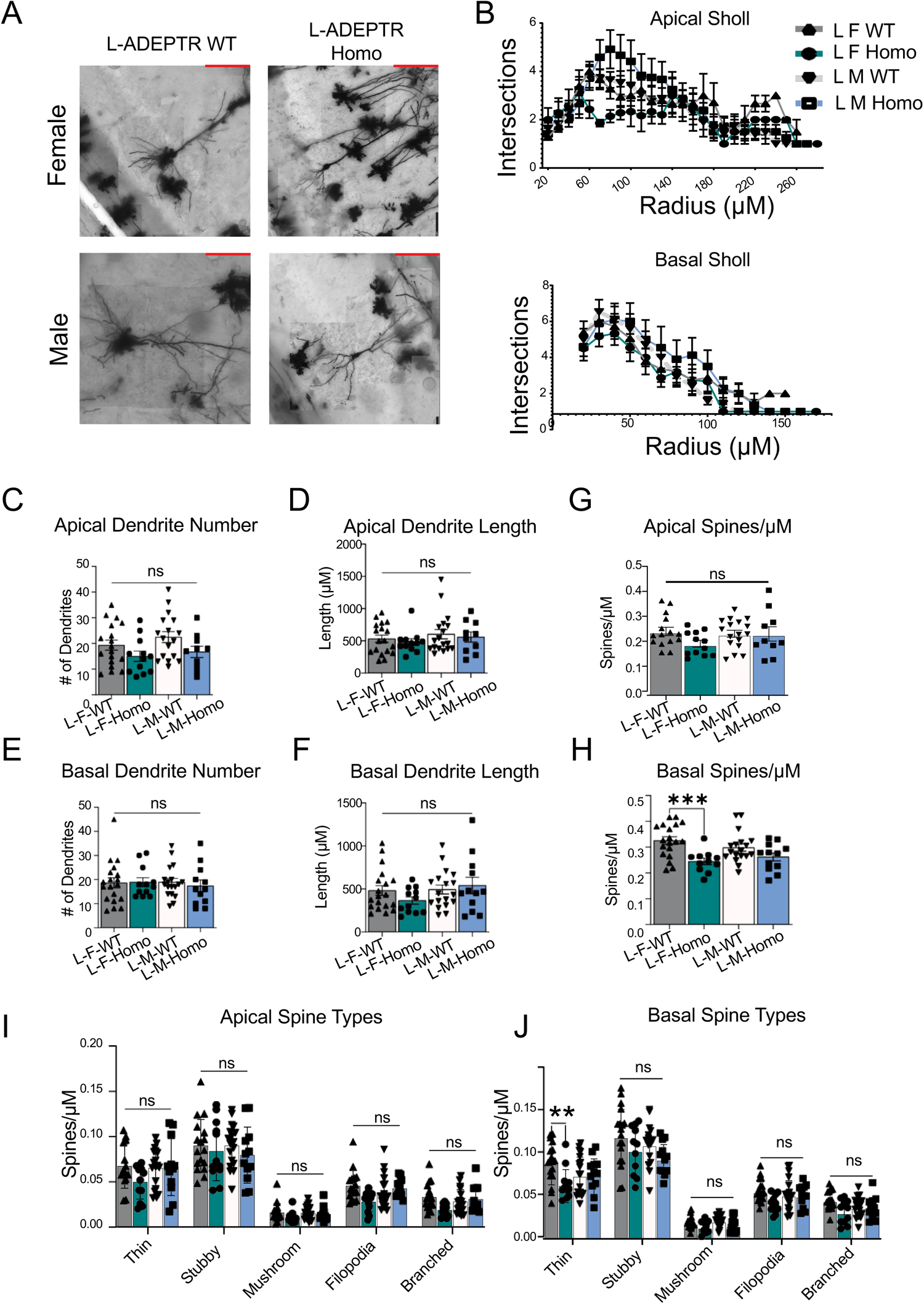
Golgi Staining of P42 L-ADEPTR CA1 pyramidal Hippocampal mouse neurons. (A) Representative images of Golgi stained CA1 pyramidal cells (male (M) and female (F) Wild-type (WT) littermate and homozygous (Homo) L-ADEPTR mice. Scale: 100uM. (B) Sholl analysis of P42 L-ADEPTR cells. (C, E) Total number of apical and basal dendrites, respectively. (D, F) Total length of apical and basal dendrites, respectively. (G, H) Spine density of apical and basal dendrites, respectively. (I, J) Spine density of different spine classes on apical and basal dendrites, respectively. (N=14-15 per group). One way ANOVA followed by Tukey’s test. Error bars indicate ± SEM. Significance level between different experimental pairs is shown (NS, not significant; \**p* < 0.05; \*\**p* < 0.01; \*\*\**p* < 0.001).

### Plasticity related gene expression is compensated in S-ADEPTR and L-ADEPTR adult mice

We next investigated whether the deficits in gene expression evident in primary neuron cultures persist into adulthood. Lack of significant changes or rescue in gene expression in the plasticity-related genes we studied in adults would suggest compensation of gene expression as animals develop and mature, thereby potentially explaining the lack of major changes in neuronal morphology and memory. Therefore, we assessed the expression of plasticity-related genes in P42 male and female S-ADEPTR and L-ADEPTR mice (Figure 10A).

**Figure 10:**
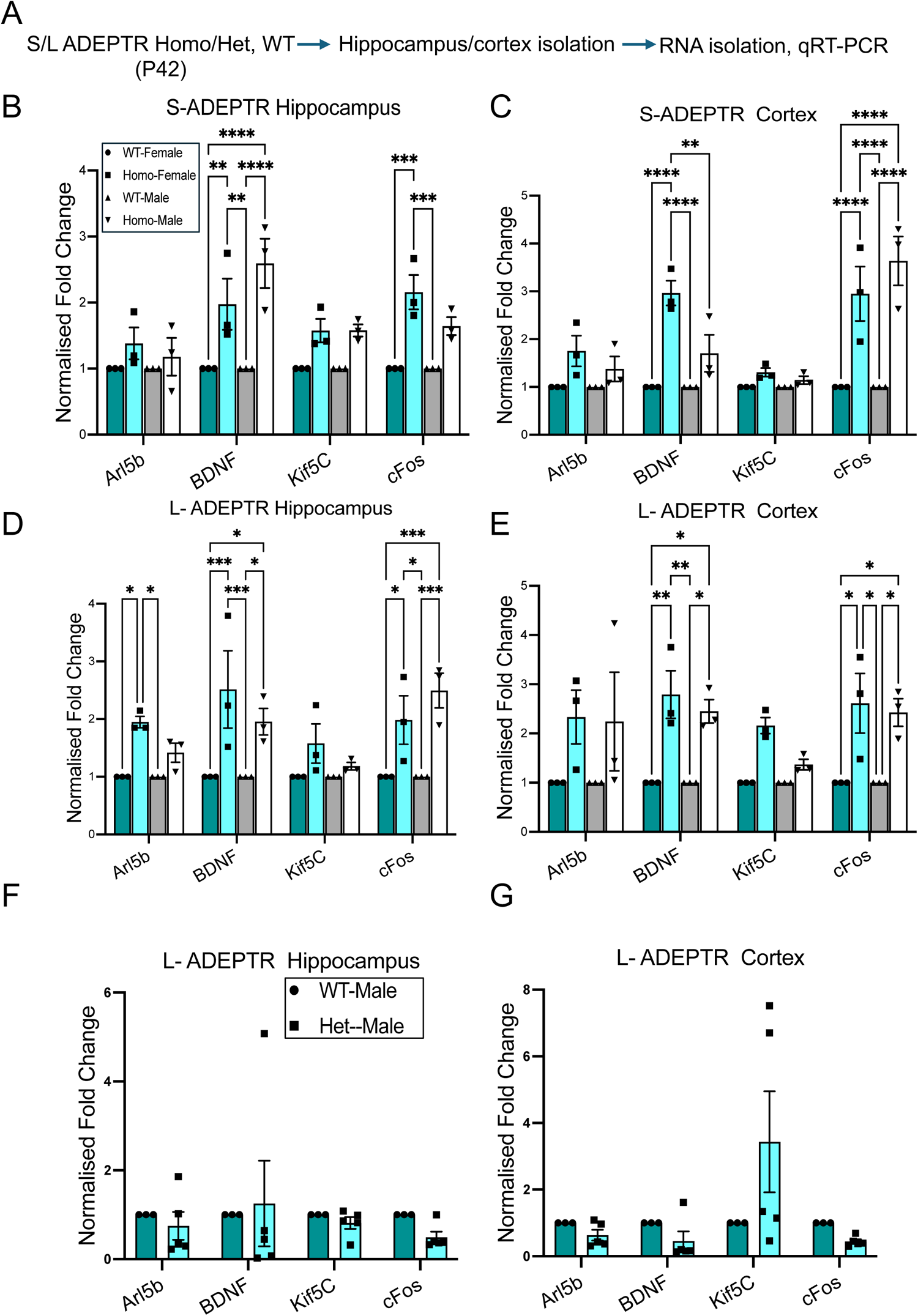
Plasticity related gene expression is enhanced in P42 SKO and LKO ADEPTR mice. (A) Experimental strategy. Isolate hippocampus and cortex from S- and L-ADEPTR P42 homozygous (Homo), Wild-type (WT) and Heterozygous (Het) mice for RNA isolation and qRT-PCR. B & C: Bar Graph showing normalized fold change of Arl5b, BDNF, Kif5C and cFos in S-ADEPTR P42 hippocampus and cortex. D & E: normalized fold change of Arl5b, BDNF, Kif5C and cFos in L-ADEPTR P42 mice hippocampus and cortex. F&G: Normalized fold change of Arl5b, BDNF, Kif5C and cFos in L-ADEPTR P42 WT and Het mice hippocampus and Cortex. N=3-4 per group), Normalization using 18S rRNA, Two way ANOVA followed by Tukey’s test for all except F & G. Paired t-test was used for F&G, Error bars indicate ± SEM. Significance level between different experimental pairs is shown (NS, not significant; \**p* < 0.05; \*\**p* < 0.01; \*\*\**p* < 0.001).

Briefly, brains of homozygous male and female P42 mice (along with age-matched male and female WT controls) were harvested, and total RNA was isolated from two brain regions, cortex, and hippocampus. Subsequently, the expression of plasticity-related genes was tested as explained before. Results showed that in S-ADEPTR, there significant changes in BDNF expression in the hippocampus of homozygous S-ADEPTR males or females compared to WT controls. Concurrently, cFos expression was elevated in homozygous S-ADEPTR females compared to WT females, but not in males. Moreover, significant differences were observed between homozygous males and females (Figure 10B, Supplementary Table S10, n= 3-4 per group, one-way ANOVA followed by Tukey’s multiple comparisons, *p < 0.05, **p < 0.005). Such differences in BDNF and cFos expression were also evident in the cortex (Figure 10C, n= 3-4 per group, one-way ANOVA followed by Tukey’s multiple comparisons, *p < 0.05, **p < 0.005). Interestingly, expression of the other two genes remained unaffected.

A similar trend of expression was evident in L-ADEPTR mice which exhibited significant changes in cFos and BDNF expression in the hippocampus of homozygous males or females compared to WT. Moreover, Arl5b expression was elevated in homozygous L-ADEPTR females compared to WT females, but not in homozygous L-ADEPTR males. (Figure 10D, n= 3-4 per group, one-way ANOVA followed by Tukey’s multiple comparisons, *p < 0.05, **p < 0.005).

Furthermore, in the cortex, there were changes in the expression of Arl5b in homozygous females compared to WT, but not males. BDNF and cFos were also exclusively elevated in homozygous females compared to WT as well as between them (Figure 10E, n= 3-4 per group, one-way ANOVA followed by Tukey’s multiple comparisons, *p < 0.05, **p < 0.005). Additionally, analysis of gene expression in P42 heterozygous male mice (L-ADEPTR) showed no significant changes when compared to WT in the hippocampus (Figure 10F) or cortex (Figure 10G).

## Discussion

In the emerging field of RNA-centric neurobiology, a major gap in knowledge is the lack of understanding of the significance, function, and mechanisms of noncoding RNAs in mediating nervous system functions. Noncoding RNAs consist of diverse classes expressed in all cells and exhibit compartment-specific enrichments. Fully understanding their in vivo functions requires developing genetic models to assess their roles at the organismal level. Several studies have shown that nuclear-enriched lncRNAs are key effectors of transcriptional changes associated with learning and memory (Spadaro et al., 2015; Butler et al., 2019; Cui et al., 2022; D. Li et al., 2018; Raveendra et al., 2018; Ben-Tov Perry et al., 2023). However, the functional and mechanistic significance of cytoplasmically enriched lncRNAs remains underexplored.

In the present study, we developed two different loss-of-function models of the lncRNA ADEPTR, which is induced by cAMP and localized to the dendrites of hippocampal neurons (Grinman et al., 2021). We conducted detailed behavioral and neuronal morphological analyses of these models, S-ADEPTR and L-ADEPTR, and found that ADEPTR loss of function results in a sex-specific phenotype.

### The significance of protein binding element of ADEPTR in governing mice anxiety-like behavior

We carried out six behavioral assessments (EPM, OF, SA, NOR, MWM, CFC/TFC) to examine deficits in anxiety-related behaviors and various types of memories in male and female mice. S-ADEPTR is characterized by a 221-base pair deletion in the ADEPTR lncRNA, which corresponds to interactions with proteins such as spectrin and ankyrin (Grinman et al., 2021). Our previous studies have shown that this fragment is sufficient to interact with spectrin and ankyrin (Grinman et al., 2021).

Our assessments revealed that male S-ADEPTR KO mice exhibit reduced anxiety, as identified from EPM data (Figure 2), whereas female S-ADEPTR and both male and female L-ADEPTR mice did not show a deficit. These observations suggest that the lncRNA-protein interaction element of S-ADEPTR is crucial for mediating anxiety-like behavior. Previously, it was shown that the lncRNA Gomafu interacts with BMI1, a key member of the polycomb repressive complex, in mediating anxiety-like behavior (Spadaro et al., 2015).

Interestingly, several lines of evidence suggest that cognition and anxiety are tightly coupled and interacting processes, given that brain regions governing anxiety, such as the hippocampus and amygdala, are also implicated in memory acquisition (Salomons et al., 2012). Since we did not identify significant deficits in any other behavioral assays, our findings underscore the significance of protein interaction elements of lncRNAs in mediating specific behavioral processes in a sex-specific manner. Although sex-specific roles of lncRNAs have been reported (Issler et al., 2020b), our findings shed new light on the sex-specific functions of lncRNAs.

### ADEPTR KOs does not affect memory

Since previous evidence showed that ADEPTR is synaptically targeted in a cAMP-dependent manner and is critical for activity-dependent structural changes at the synapse, we anticipated that ADEPTR loss-of-function might impact memory formation in adult mice. However, contrary to our expectations, both S-ADEPTR and L-ADEPTR KO models showed no deficits in NOR, MWM, SA, or CFC/TFC tasks (Figures 2, 3, Supplementary Figures 1-4).

Previous studies on ADEPTR function utilized shRNA-mediated knockdown, which resulted in only partial loss of function but produced significant deficits in excitatory synaptic transmission and cAMP-dependent changes in synapse density and morphology. Additionally, knockout studies on a key nuclear-enriched lncRNA, Malat1, revealed that Malat1 is dispensable for mouse development (Zhang et al., 2012; Eißmann et al., 2012). Taken together, these observations suggest the existence of compensatory mechanisms in vivo that mitigate memory deficits.

### Systematic Assessments of neuronal morphology reveal early deficits that are compensated in a sex specific manner in S and L-ADEPTR KOs

In search of a putative compensatory mechanism, we first investigated whether the deficits observed with shRNA-mediated knockdown of ADEPTR in hippocampal neurons would also be present in ADEPTR knockout models. We cultured neurons from male and female S-ADEPTR and L-ADEPTR pups (postnatal days 1-3). Consistent with our earlier findings (Grinman et al., 2021), we observed morphological deficits in the ADEPTR KOs. Interestingly, these deficits were more severe than previously reported, with reductions in dendritic arborization, spine density, and morphology in both male and female S-ADEPTR and L-ADEPTR pups. These results suggest that compensation is not occurring prenatally.

To further investigate, we examined neuronal morphology in two additional age groups: 14 days old and 42 days old male and female S-ADEPTR and L-ADEPTR mice. Golgi staining followed by analysis of apical and basal dendrite number, length, density, and branching in CA1 pyramidal neurons of the hippocampus indicated that by postnatal day 14 (P14), all the deficits observed in neurons from postnatal days 1-3 (P1-3) were compensated in both male and female S-ADEPTR and L-ADEPTR mice. However, analysis of P42 neurons identified a deficit in thin spines in female L-ADEPTR mice without any changes in the other parameters measured in both males and females (Figures 8 and 9).

These assessments identified a critical time window (postnatal day 3 to postnatal day 14) during which the morphological deficits are compensated. However, morphological deficits begin to reappear in P42 L-ADEPTR females. These observations suggest that these KO models might exhibit deficits at later ages.

### Analyses of Plasticity related gene expression suggest possible compensatory mechanisms

Drawing parallels from the Malat1 KD study (Zhang et al., 2012; Eißmann et al., 2012), we posited that the absence of a strong memory deficit phenotype in adult ADEPTR KO mice might be due to the recovery of plasticity-related gene expression along with normal neuronal morphology. To validate this, we assayed the hippocampus and cortex of P42 male and female mice for the expression of BDNF, a known plasticity-related gene (reviewed in Miranda et al., 2019); cFos, an immediate early gene (reviewed in Minatohara et al., 2016); Kif5C, a kinesin critical for neuronal morphology and memory (Swarnkar et al., 2021); and Arl5b, the host gene of ADEPTR (Grinman et al., 2021).

Our qPCR results (Figure 10) show a significant increase in BDNF expression in both male and female S-ADEPTR KO mice, along with a consistent increase in cFos expression. In female L-ADEPTR KO mice, we observed consistent enhancements in the expression of Arl5b, BDNF, and cFos, while their male counterparts showed consistent increases in BDNF and cFos levels. These results suggest that the activation of several plasticity-related genes in these KOs might underlie the lack of memory deficits in both S-ADEPTR and L-ADEPTR KO male and female mice.

However, the lack of memory deficits could be more complex than just the enhancements in the expression of these genes. For instance, while enhancements in Kif5C in the dorsal CA1 were sufficient to improve MWM learning, they did not affect performance in contextual fear conditioning, indicating memory-specific effects of Kif5C overexpression. Similarly, BDNF overexpression has been shown to produce learning impairments (Cunha et al., 2009). Taken together, our results indicate that genomic deletion of ADEPTR, although it results in impairments in neuronal morphology immediately after birth, leads to specific transcriptional changes later in development that ameliorate these deficits. Our observations suggest that enhancements in the expression of plasticity-related genes are a component of this transcriptional change.

In summary, the loss-of-function models we developed (S-ADEPTR and L-ADEPTR KOs) reveal the intricate roles that lncRNAs and their regulatory elements may have in the nervous system. The male-specific anxiety-like behavior deficit in S-ADEPTR KO mice suggests that although the loss of function induces long-term transcriptional changes, these changes do not entirely restore normalcy. Importantly, our findings highlights the sex-specific role of the protein-interacting region of ADEPTR in mediating anxiety-like behavior. In the L-ADEPTR model, neither males nor females showed impairments in various behavioral tests, including anxiety-like behavior. However, females had a decreased number of thin spines in the basal dendrites, further indicating sex-specific functions of ADEPTR. Investigating the sex-specific roles of ADEPTR and the mechanisms behind the observed compensations will provide valuable insights into the complex regulation and functions of lncRNAs.

## Materials and Methods

### Animals

#### Generation of ADEPTR knockout (KO) mice

We generated ADEPTR KO mice using CRISPR-Cas9 based on previously reported methods (Nishizono et al., 2020). Briefly, 100 ng/uL of gRNA (produced by annealing crRNA and tracrRNA) and 10 ng/uL of Cas9 protein were introduced into C57BL/6N mouse fertilized eggs using a NEPA21 electroporator (Nepa Gene Co., Ltd., Ichikawa, Japan). crRNA, tracrRNA, and Cas9 were all purchased from IDT. The day after electroporation, fertilized eggs that developed to the 2-cell stage were embryo transferred to pseudo-pregnant mice to generate F0 mice. Primer sets for genotyping are included in Supplementary Table S1.

#### Breeding Scheme

Breeding schemes established for both strains, regarding behavioral trials, consisted of heterozygous males crossed with heterozygous females. Breeding schemes established for both strains, regarding cell culture and Golgi straining, consisted of homozygous males crossed with homozygous females; as well as WT males crossed with WT females. The S-ADEPTR strain had minimal difficulties in breeding performance. To produce adequate mice for behavioral experiments, several monogamous breeding pairs were established. Each cage included several forms of enrichment: a mouse hut and two nestlets. Immediately after the doe was identified as pregnant, the buck was removed, and the doe remained singly housed until offspring were weaned. All monogamous breeding pairs were set up simultaneously to ensure offsprings met the parameters of the behavior experiment.

Whilst breeding the L-ADEPTR strain, we noticed a decline in breeding performance. Mice were producing very small litters and not providing essential care leading to deaths of entire litters. To produce adequate mice for behavioral experiments, several breeding trios were established. Each cage included several forms of enrichment: a mouse hut and two nestlets. Once each doe was identified as pregnant, the buck was removed, and each mouse remained singly housed until offspring were weaned. All breeding trios were set up simultaneously to ensure offspring met the parameters of the behavior experiment. Mice were maintained on a 12hr light/dark cycle with ad libitum access to water and food. In vivo experiments were carried out during the light part of the cycle light/dark cycle. All in vitro experiments were performed in primary hippocampal cell cultures obtained from C57BL6 mouse pups. Housing and experimental procedures were approved and supervised by the Institutional Animal Care and Use Committee of the Herbert Wertheim UF Scripps Institute for Biomedical Innovation & Technology and Max Planck Florida Institute for Neuroscience.

### Behavior

For behavioral assays, all mice are handled for 3 days prior to commencement of testing. The experimenter was blind to group conditions during testing. Test order was determined randomly (or pseudo-randomly, with random assignment and verification that each group was evenly distributed across “x factors”). Approximately 70dB background white noise was used during all tests except for water maze testing. At least 30 minutes prior to testing on any day, mice were moved from their holding room to a waiting room near the experimental assay room where they were allowed to rest prior to being tested, except where noted below.

#### Elevated Plus Maze

An elevated plus maze test was used to assess baseline anxiety-like behavior. Mice were placed in the center of the plus maze (black maze with white flooring; Med Associates, St. Albans, VT) and were allowed to explore the maze for 5 minutes. Time spent in open and closed arms, number of arm entries, latency to initially enter an open arm and total distance moved were recorded using EthoVision XT (Noldus Information Technology Inc., Leesburg, VA). Uniformity in lighting was confirmed across the maze, and the maze was cleaned with Quatracide PV before each trial. Lighting for the maze was set at 200 lux in the center of the plus maze, 270 lux on the open arms, and 120 lux on the closed arms.

#### Open Field

To test for baseline activity, locomotor behavior was measured in 17×17 in. square acrylic open field chambers. Prior to testing, uniformity of light across the arena was confirmed using a light intensity meter, and the chambers were cleaned with Quatracide PV before and between trials. Mice were placed into the center of the chamber to begin testing, and activity was recorded for 30 min using EthoVision XT (Noldus Information Technology Inc., Leesburg, VA). Data were analyzed in 10 min blocks for activity and in one 5 min block for anxiety-like behavior (thigmotaxis).

#### Spontaneous Alternation

Working memory was assessed in a spontaneous alternation T-maze test. Mice received 2 tests, each separated by one week. Two mazes were used, each turned in a different direction, to increase novelty to the maze on the second test. Each maze (Med Associates, St. Albans, VT) contained three arms with walls that blocked visual interruption and included a start box at the base of the start arm. Automatic guillotine doors were installed at the entry of each arm that were controlled by EthoVision XT (Noldus Information Technology Inc., Leesburg, VA). Each test was conducted thusly: a mouse was placed in a start box and the door to the maze subsequently opened, allowing the mouse to enter the maze and explore to the T intersection. Upon reaching the intersection, the mouse chose an arm (free choice trial) and, after 3 body points had entered that arm, the door closed automatically, detaining the mouse in that arm for a period of 10 seconds. During those 10 seconds, a cloth lightly sprayed with 70% ethanol was used to wipe the maze floor outside the chosen arm to remove possible odor cues. After 10 seconds, the mouse was placed back in the start box for a second free choice trial, after which the door to that arm again closed, detaining the mouse in that arm until prompt removal. Of the 2 tests the mouse was given, the first kept the mouse in the start box for 60s before the door opened to allow the mouse to enter the maze for a second trial (delay condition), while the second test allowed the mouse to immediately enter the maze for the second trial (no delay condition). Groups were balanced for maze and test day. Alternation success was calculated for each test. If a mouse did not leave the start box to enter the maze after 45s, it was gently nudged with forceps. The maze was cleaned with 70% ethanol between mice.

#### Novel Object Recognition

Non-spatial memory was assessed using the novel object recognition (NOR) test. On days 1 and 2, mice were habituated to the NOR arena (two 17×17 inch white open fields with top-mounted internal lighting) for 10 minutes each day. On day 3, a set of 2 identical objects (2 sets available that differed in color and texture) were placed into the back corners of each maze. Each object was permanently affixed to a disk that fit into the floor of the maze to ensure objects could not be moved once placed and that object placement was identical in each maze. Mice were placed into the front center of each maze and given a maximum of 20 minutes to investigate the objects. Behavior was recorded using EthoVision XT with Deep Learning, and object investigation was automatically monitored until mice accumulated a total of 60 seconds of investigation of the objects. Once this criterion was met for each mouse, the trial ended, and the mouse was immediately returned to its cage. On day 4, one object from each arena was swapped with an object from the other arena, thus creating a “novel” object. Mice were again placed into the front center of each maze and were given 10 minutes to investigate the objects. Total time spent investigating each object was recorded. The arenas were cleaned with Quatracide PV and the objects were cleaned with 70% ethanol prior to each trial. Investigation was defined as the head being directed toward any of an 8cm zone around the object while the nose was located within a 12cm zone around the object. This definition included periods when the mouse was physically interacting with the object. A preference score was calculated for each mouse as duration (s) exploring the novel object ÷ (duration (s) exploring novel + duration (s) exploring familiar).

#### Morris Water Maze

The Morris Water Maze (MWM) was used to test for spatial memory and cognitive flexibility. The test was performed in a 1.4 m diameter white tank with a 10 cm diameter platform submerged approximately 1 cm below the surface of the water. The water was made opaque using non-toxic white washable paint that made the platform invisible during trials. The temperature of the water was kept at 22°C. A curtain surrounded most of the tank and visual cues were placed on the curtain for spatial reference. Prior to water maze training, mice received a visual platform test where the spatial cues were removed, and the platform was elevated above the surface of the water and marked with a cue so it could clearly be discerned. Mice were given four trials, and the platform location was varied over trials. This served to habituate the mice to the task, to verify the visual ability of the mice and to ensure that the mice had no deficits that would affect their ability to swim to the platform. For the water maze spatial learning paradigm, mice were given four acquisition trials per day until they reached a pre-determined criteria of an average latency (by group) of 20s to find the platform and at least 95% success in finding the platform. Activity and performance were tracked using EthoVision XT (Noldus Information Technology, Leesburg, VA). The start location was varied for each trial, and mice were allowed 60 seconds to find the platform. Mice were left on the platform for 15 seconds before removing them from the water maze. If a mouse did not find the platform within 60 seconds, it was placed on or guided to the platform and kept there for 15 seconds. Mice were dried after each trial and placed into cages located atop heating pads to prevent hypothermia. Daily acquisition trials were averaged for analysis. To verify learning progress, after every 3 days of training, mice were given a probe trial during which the platform was not present, and a novel start location was used for these trials. These probe trials were given the morning after each 3^rd^ day of acquisition training prior to commencement of the next day’s training. 24 hours after reaching criteria, mice were given a final probe trial. Total time spent in each quadrant, total number of platform crossings, latency to first platform crossing, and average distance to the platform center were recorded for a 60 second probe trial.

Following the final learning probe trial, mice were trained in a reversal learning paradigm where the platform was moved to the opposite quadrant location as before and mice were trained to find the platform in this new location. Again, mice were trained until they reached criteria. Mice received probe trials throughout the reversal learning phase of the test every 3 days as in the learning paradigm and then again at 24 hours after reaching criteria for the new platform location. The same dependent variables were collected.

#### Contextual Fear Conditioning

The fear conditioning chambers (Ugo Basile) consisted of 26×26×35H cm Plexiglas chambers with top-mounted cameras, housed inside sound-attenuating chambers. Infrared lighting was utilized for all behavior tracking using EthoVision XT software (Noldus Information Technology, Inc., Leesburg, VA). Each chamber was cleaned with 70% ethanol prior to each trial. White light was used inside the chamber for training and testing, and white noise (approximately 70dB) was played in the room to mask any unintended noise that might add to the context. The tray below the shock grid floor was lined with Techboard Ultra paper (Shepherd Specialty Papers) and was changed out after each trial. The back wall of each chamber was lined with a black and white checkerboard pattern. Mice were placed individually into each chamber and allowed to explore it for 2.5 min. Mice then received either one or two (with a 60s ITI) two-second 0.5 mA foot shocks. 30 s after, the mice were removed from the chambers and returned to their home cages. 24 h later, mice were tested for contextual fear conditioning by placing them back into the chambers for 5 min. Freezing, defined as the absence of movement except that required for breathing, was determined using the immobility variable in EthoVision XT(Pham et al., 2009), and a subset of trials was analyzed by a blind observer for accuracy of detection. Percent of time spent freezing and activity (expressed as total distance moved) were recorded. Mice received an additional context test (performed exactly as the first one) one-week post-training, and again percent freezing, and activity were recorded.

#### Trace Fear Conditioning

The fear conditioning chambers (Ugo Basile) consisted of 26×26×35H cm Plexiglas chambers with top-mounted cameras, housed inside sound-attenuating chambers. Infrared lighting was utilized for all behavior tracking using EthoVision XT software (Noldus Information Technology, Inc., Leesburg, VA). Training and context testing occurred in context A and trace testing occurred in context B. Context A consisted of the fear conditioning chambers with a black and white checkerboard patterned back wall and white chamber lighting. 70dB white noise was played in the testing room during training and room lights were on. The chamber was cleaned with 70% EtOH. The tray below the shock grid floor was lined with Techboard Ultra paper (Shepherd Specialty Papers) and was changed out after each trial. On day 1, mice were placed individually into each chamber and allowed to explore it for 3 min. Mice then received 5 presentations of 20s tone (2700hz 80dB), 20s delay, and a 2s shock (0.5mA), each with a 200s ITI. 60s following the last shock, mice were removed from the chambers. On day 2, mice were brought directly from the holding room to the testing room before being tested for trace fear conditioning in context B. In context B, the chambers were disguised with white inserts that changed the look, dimensions and feel of the chamber, and with orange extract placed inside the sound attenuating cubicles to change the smell of the chamber. White light was used inside the chamber. 70% isopropanol was used instead of ethanol to clean the chambers, as well, and white noise and light was not used in the testing room during the trials. Following a 2-min habituation, mice received 3 presentations of the same 20s tone as on day 1, each with a 220s ITI. Mice were removed from the chambers immediately following the last tone. On day 3, mice were tested for contextual fear conditioning by placing them back into context A for 5 minutes. Freezing, defined as the absence of movement except that required for breathing, was determined using the immobility variable in EthoVision XT(Pham et al., 2009), and a subset of trials was analyzed by a blind observer for accuracy of detection. Percent of time spent freezing and activity (expressed as total distance moved) were recorded on all days.

### Quantitative real-time PCR (qRT-PCR)

The RNA from the cell cultures, hippocampal or cortical tissues were extracted with TRIzol reagent and purified with Zymo Direct-Zol RNA Microprep kit. RNA was reverse transcribed to cDNA using the same method previously reported from this laboratory (Kadakkuzha et al., 2013;Raveendra et al., 2018). 1µg of RNA was used with Quanta cDNA SuperMix (Quanta Biosciences, Gaithersburg, MD) according to the manufacturer’s instructions and the expression of transcripts were quantified by qRT-PCR using SYBR Green PCR master mix (Applied Biosystems Carlsbad, CA) for detection in QuantStudio 6 Pro cycler (Applied Biosystems Carlsbad, CA). Quantification of each transcript was normalized to the mouse 18S reference gene following the 2^-ΔΔCt^ method ((Livak & Schmittgen, 2001); Kadakkuzha et al., 2013). One-way ANOVA, Two Way ANOVA followed by Tukey’s Test or Paired t test were used to identify genes with statistically significant expression levels. Primer sets for qPCR are included in Supplementary Table S1.

### Primary Hippocampal Cell Cultures

Neuronal cultures from hippocampus were obtained from brains of P1-P3 mice. Briefly, brains were removed from the skulls and membranes cleaned. Then, brains were dissociated using papain (29.5 U/mg protein, Worthington). The cells obtained were plated in poly-D-lysine (PDL) treated plates with a 5×10^5^ density (for RNA) or a density of 1×10^5^ in 24 well black confocal plates (for imaging). Both types of cultures were maintained in Neurobasal medium (Invitrogen), penicillin/streptomycin and 2% B27 (Invitrogen) at 37°C in 5% CO2.

### Constructs & Transfections

For lipofectamine transfection, about 1.50 × 10^5^ cells/well primary mouse hippocampal neurons were plated on PDL-coated CellVis glass-bottom 24-well plate (P24-1.5H-N). For Negative Control-GFP arborization analysis, neurons were transfected 68-74 hrs. prior to the imaging (DIV 14) using Lipofectamine 2000 (Thermo) following manufacturer’s instruction. Before adding the DNA-lipofectamine mixture, half of the conditioned culture medium was removed and saved for later. Four hours after incubation with DNA-lipofectamine mixture, the medium was removed and 500ul of 1:1 of conditioned/fresh medium was immediately added. Right before imaging, all culture medium was replaced by 500 ul of Hibernate E low fluorescence buffer (BrainBits) to maintain the ambient pH environment.

### Morphology Assessments

After 72 hrs. of transfection, mice hippocampal neurons expressing NC-GFP, images were collected at 36°C in the light microscopy facility at UF Scripps Biomedical Research, using a confocal microscope (FV1000; Olympus; Apo N 60X/1.49 Oil) in Hibernate-E (Brainbits). Z-stack images were acquired using Fluoview1000 (64 bit) software (Olympus) and converted into a maximum projection intensity image in FIJI (ImageJ, NIH). Dendritic arbor was quantified via the Sholl analysis plugin in FIJI. The center of soma was considered as the midpoint and origin of the concentric radii was set from that point to the longest axis of soma. The parameters set for analysis were starting radius 20 μm, ending radius 100 μm, radius step size 10 μm. The maximum value of sampled intersections reflecting the highest number of processes/branches in the arbor was calculated and the number of intersections plotted against distance from the soma center in μm. Data was analyzed using Two-way ANOVA followed by Tukey’s test.

Spine morphology was analyzed using MATLAB software developed in the light microscopy facility at the Max Planck Florida Institute. By using a geometric approach, this software automatically detects and quantifies the structure of dendritic spines from the selected secondary branch (100 μm length) in the Z-stack confocal image. The software assigns the detected spines to one of the three morphological categories (thin, stubby or mushroom) based on the difference in structural components of the spines i.e., head, neck and shaft. Student’s t test was carried out to evaluate the statistical difference amongst the groups.

### Golgi Cox staining

Mice brains were harvested at either p14 or p42, stored in 4% PFA in PBS in 4 degrees overnight. Next day they were transferred to Golgi-cox solution (Solution 1-potassium dichromate and mercuric chloride in warm DI water, Solution 2-potassium chromate in cold DI water, slowly poured second solution into first solution, stored in dark for minimum of 72 hours. Used orange supernatant fluid from this for Golgi staining), stored for 14 days in dark. Next, they were transferred to 30% sucrose in PBS overnight, or until brains sunk to bottom. Brains were rinsed and pat dried, and frozen in cryo OCT in −80.

Processed brains were sectioned at 80uM in Leica cryostat at −20 degrees, mounted on gel coated coverslip. For dehydration, brain slices were treated with ddH2O for 5 mins, 20% ammonium hydroxide for 10 mins, ddH2O for 5 mins, 70% EtOH for 30 seconds, 95% EtOH for 30 seconds, 100% EtOH for 30 seconds, then Xylene for 40 mins. Finally, sections were mounted on glass coverslips and sealed with nail polish. Sections were imaged on Nikon e800 microscope at both 20x and 60x using Neurolucida software.

#### N values

<u>P14:</u> S-F-WT: 14 cells, 3 brains; S-F-Homo: 15 cells, 3 brains; S-M-WT: 19 cells, 3 brains; S-M-Homo: 16 cells, 3 brains; L-F-WT: 17 cells, 3 brains; L-F-Homo: 15 cells, 3 brains; L-M-WT: 17 cells, 3 brains; L-M-Homo: 16 cells, 3 brains

<u>P42:</u> S-F-WT: 18 cells, 3 brains; S-F-HOMO: 12 cells, 3 brains; S-M-WT: 18 cells, 3 brains; S-M-HOMO: 12 cells, 2 brains; L-F-WT: 18 cells, 3 brains; L-F-Homo: 12 cells, 3 brains; L-M-WT: 18 cells, 3 brains; L-M-Homo: 12 cells, 2 brains.

### Statistics

All statistical analysis was performed in Excel and GraphPad Prism 10. Data are represented by the mean, and error bars represent SEM. Statistical tests performed are unpaired two-tailed *t* test, one-way and two-way ANOVA with Tukey’s post hoc test, Dunnett’s test, or pairwise *t* test, as indicated. *N* represents the number of biological replicates for each experiment, unless stated otherwise. Outliers were identified and excluded using the ROUT method in GraphPad prism version 10, with a Q=1%.

## Data Availability

The current manuscript describes generation of knockout mouse models and detailed characterization, so no data have been generated for this manuscript. All data used for generating graphs in the Main and Supplementary Figures are included in the supplementary Tables.

## Supplemental Information

There are four supplementary Figures and ten supplementary tables accompanying this manuscript.

## Acknowledgements

We gratefully acknowledge funding support from NIH (1R01MH119541-01A1, R21 MH127734 and 5R21DA039417-02), and a Training Grant in Alzheimer’s Drug Discovery from the Lottie French Lewis Fund of the Community Foundation for Palm Beach and Martin Counties, Florida, enabling us to carry out this work. We are thankful to Ms. Erin Bryant for help with genotyping and initiating mouse behavior studies, Dr. Damon Page of UF Scripps for helpful discussion regarding Golgi staining and Dr. Baoji Xu of UF Scripps for the help with imaging of Golgi stained brain slices.

## Author Contributions

EG,HN,RY and SVP conceptualized the study and initiated the project. HN and RY generated knockout models with input from EG and SVP. EG and IE established the mouse lines with the help of GW and AP. EG and BLR developed genotyping strategies, BLR and SL carried out genotyping. GW and AP maintained the mouse lines and carried out backcrossing. AB carried out all behavior assessments. KC carried out imaging in primary neuronal cultures, and molecular analyses, JPC carried out Golgi staining and analyses. JW facilitated maintaining mouse lines, imaging and analyses, KC and SVP wrote the paper with inputs from all authors.

## Supplementary Figures

**Supplementary Figure 1.**
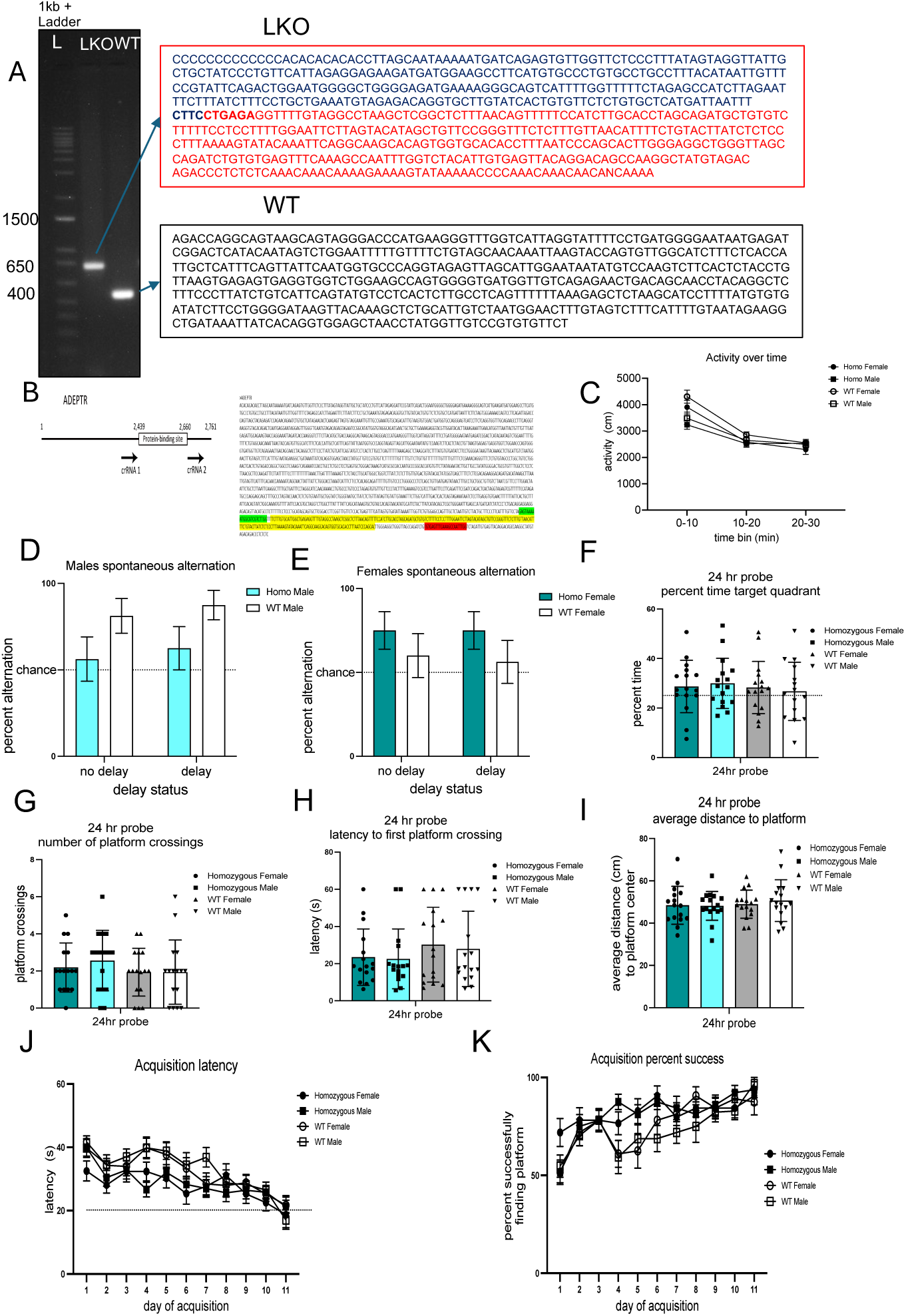
Characterization of ADEPTR knockout (KO) mice. (A) Gel analysis of PCR products from LKO mouse. Presence of the band at 650bp with LKO primers shows the mouse is HOMO. WT primers don’t give any band for HOMO samples. A band around 500bp with WT primers indicate WT. LKO primer doesn’t give any amplification with WT samples as the amplicon is too big to yield a band at the conditions we used. For HET samples both primers gave the PCR amplification band. (B) Diagram of crRNA designs and sequence of ADEPTR. Yellow highlights indicate protein binding sites. Green and red highlights indicate crRNAs, respectively. (C) Line Graph depicting activity (cm) vs time in Open Field Test of indicated phenotypes (N=14-15 per group) of S-ADEPTR mice. (D) and (E) Bar Graph depicting percent alternation in spontaneous Alternation of indicated phenotypes (N=14-15 per group) of S-ADEPTR mice. (F) Percent time spent in target quadrant of Water Maze of indicated phenotypes (N=14-15 per group) of S-ADEPTR mice. (G) Platform crossings in Water Maze of indicated phenotypes (N=14-15 per group) of S-ADEPTR mice. (H) Latency (s) for first platform crossing in Water Maze of indicated phenotypes (N=14-15 per group) of S-ADEPTR mice. (I) Average distance to platform center in Water Maze of indicated phenotypes (N=14-15 per group) of S-ADEPTR mice. (J) and (K) Acquisition latency and percent success in Water Maze of indicated phenotypes (N=14-15 per group) of S-ADEPTR mice. One way ANOVA followed by Tukey’s test. Error bars indicate ± SEM. Significance level between different experimental pairs is shown (NS, not significant; \**p* < 0.05; \*\**p* < 0.01; \*\*\**p* < 0.001).

**Supplementary Figure 2.**
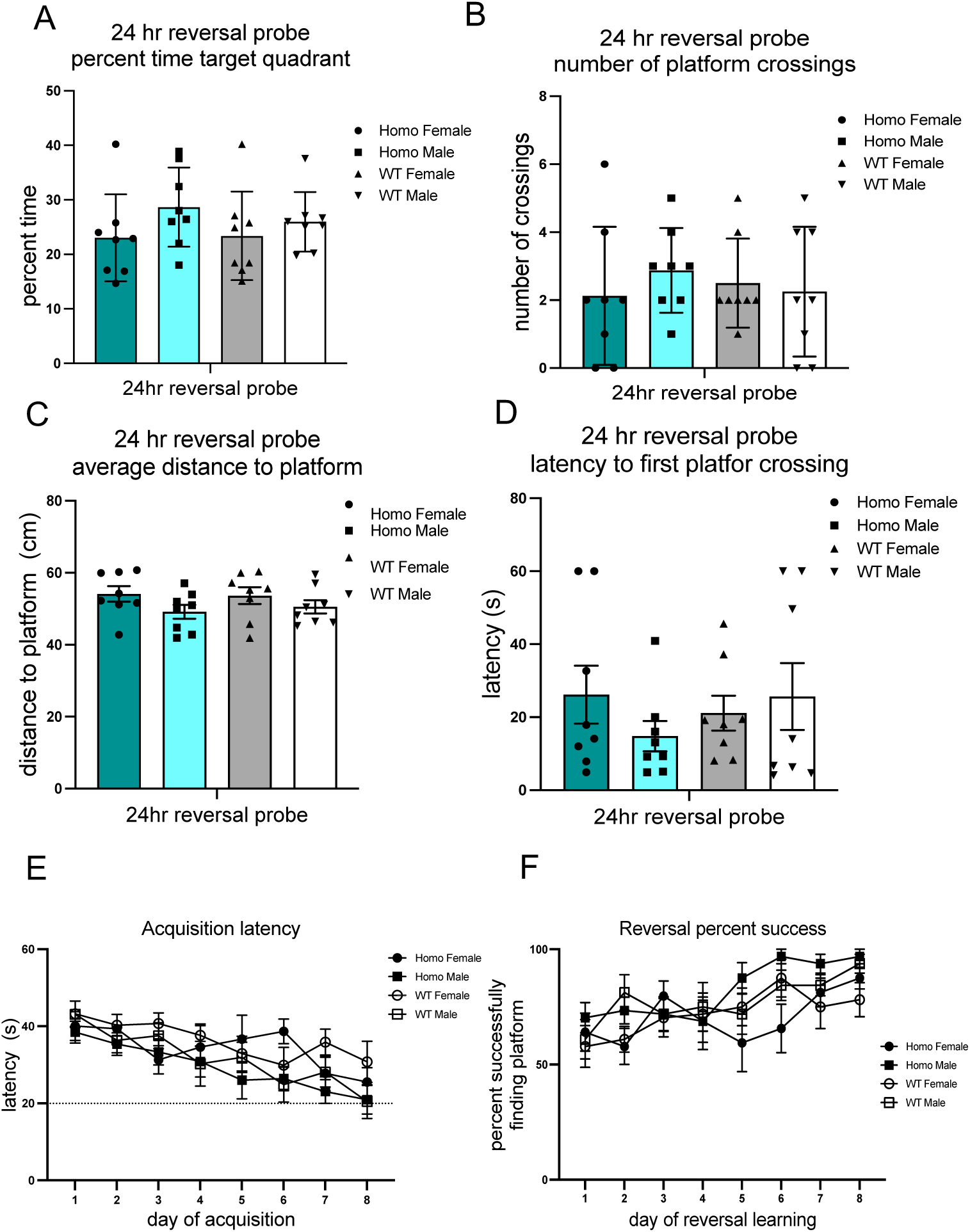
Reversal learning in MWM of S-ADEPTR KO mice. (A) Bar Graph depicting percent time spent in target quadrant of Water Maze of indicated phenotypes (N=14-15 per group) of S-ADEPTR mice. (B) Platform crossings in Water Maze of indicated phenotypes (N=14-15 per group) of S-ADEPTR mice. (C) Average distance to platform center in Water Maze of indicated phenotypes (N=14-15 per group) of S-ADEPTR mice. (D) Latency (s) for first platform crossing in Water Maze of indicated phenotypes (N=14-15 per group) of S-ADEPTR mice. (E) and (F) Acquisition latency and percent success in Water Maze of indicated phenotypes (N=14-15 per group) of S-ADEPTR mice. One way ANOVA followed by Tukey’s test. Error bars indicate ± SEM. Significance level between different experimental pairs is shown (NS, not significant; \**p* < 0.05; \*\**p* < 0.01; \*\*\**p* < 0.001).

**Supplementary Figure 3.**
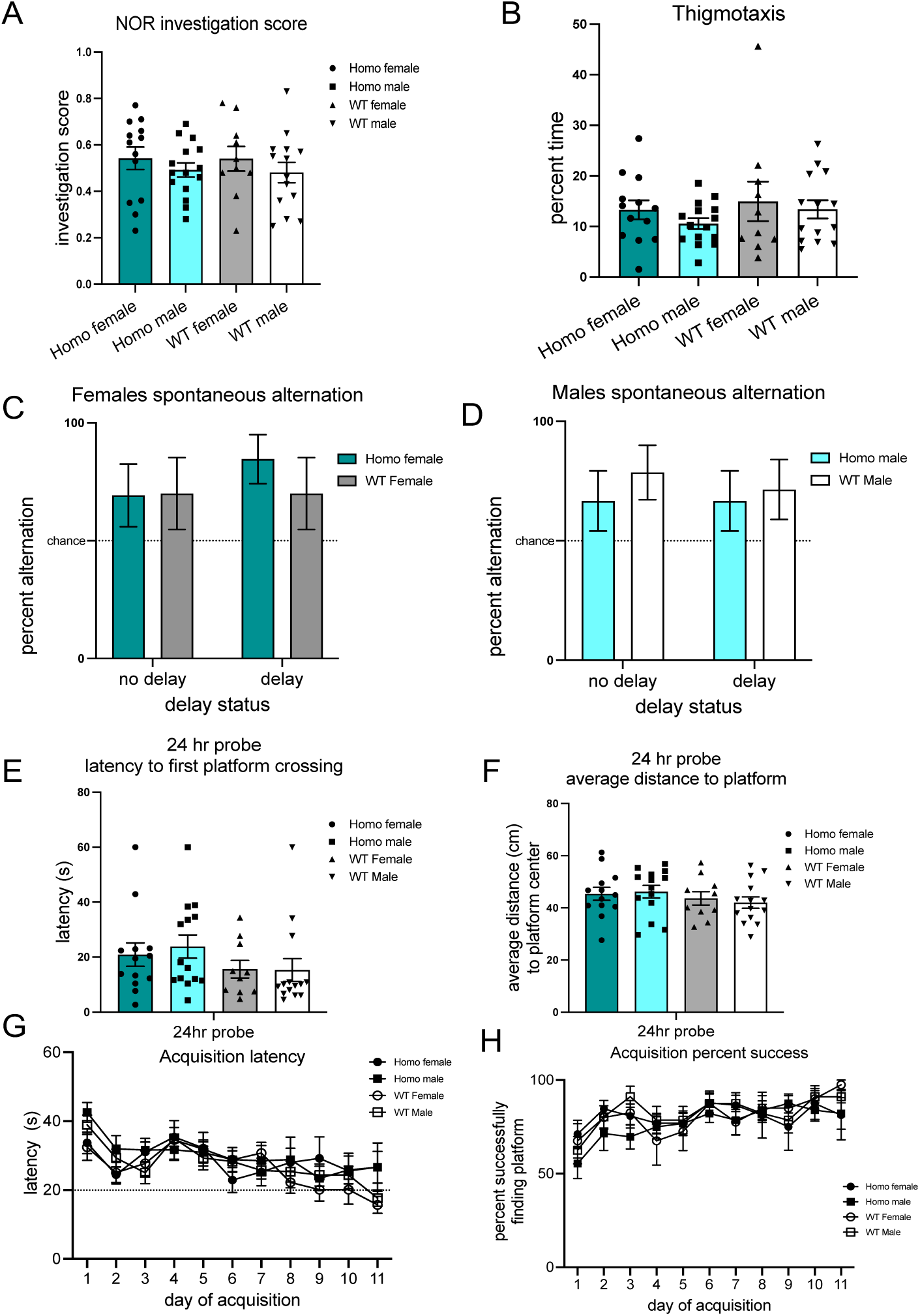
NOR, OF, SA and MWM of L-ADEPTR KO mice. (A) Bar Graph depicting investigation score of Novel object recognition in indicated phenotypes (N=15-16 per group) of L-ADEPTR mice. (B) Percent time in Thigmotaxis of indicated phenotypes (N=15-16 per group) of L-ADEPTR mice. (C) and (D) Percent alternation in spontaneous Alternation of indicated phenotypes (N=14-15 per group) of L-ADEPTR mice. (E) Latency (s) for first platform crossing in Water Maze of indicated phenotypes (N=14-15 per group) of L-ADEPTR mice. (F) Average distance to platform center in Water Maze of indicated phenotypes (N=14-15 per group) of L-ADEPTR mice. (G) and (H) Line Graph depicting acquisition latency and percent success in Water Maze of indicated phenotypes (N=14-15 per group) of L-ADEPTR mice. One way ANOVA followed by Tukey’s test. Error bars indicate ± SEM. Significance level between different experimental pairs is shown (NS, not significant; \**p* < 0.05; \*\**p* < 0.01; \*\*\**p* < 0.001).

**Supplementary Figure 4.**
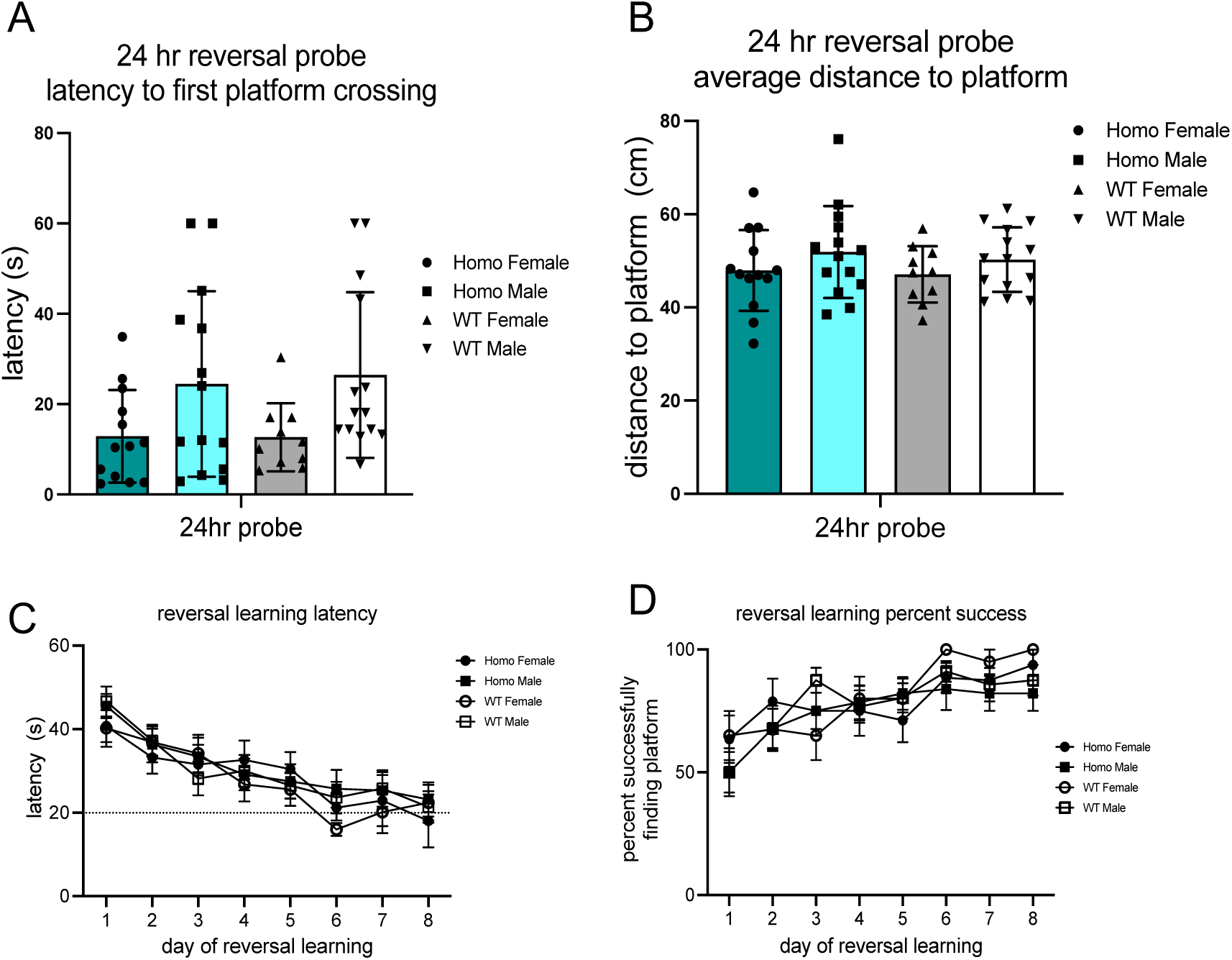
Reversal learning in MWM of L-ADEPTR KO mice. (A) Bar Graph depicting platform crossings in Water Maze of indicated phenotypes (N=14-15 per group) of L-ADEPTR mice. (B) Average distance to platform center in Water Maze of indicated phenotypes (N=14-15 per group) of L-ADEPTR mice. (C) and (D) Line Graph depicting acquisition latency and percent success in Water Maze of indicated phenotypes (N=14-15 per group) of L-ADEPTR mice. One way ANOVA followed by Tukey’s test. Error bars indicate ± SEM. Significance level between different experimental pairs is shown (NS, not significant; \**p* < 0.05; \*\**p* < 0.01; \*\*\**p* < 0.001).

